# Multi-model preclinical platform predicts clinical response of melanoma to immunotherapy

**DOI:** 10.1101/727826

**Authors:** Eva Pérez-Guijarro, Howard H. Yang, Romina E. Araya, Rajaa El Meskini, Helen T. Michael, Suman Kumar Vodnala, Kerrie L. Marie, Cari Graff-Cherry, Sung Chin, Anthony J. Iacovelli, Alan Kulaga, Anyen Fon, Aleksandra M. Michalowski, Willy Hugo, Roger S. Lo, Nicholas P. Restifo, Terry Van Dyke, Shyam K. Sharan, Romina S. Goldszmid, Zoe Weaver Ohler, Maxwell P. Lee, Chi-Ping Day, Glenn Merlino

## Abstract

Although immunotherapy has revolutionized cancer treatment, only a subset of patients demonstrates durable clinical benefit. Definitive predictive biomarkers and targets to overcome resistance remain unidentified, underscoring the urgency to develop reliable immunocompetent models for mechanistic assessment. Here we characterize a panel of syngeneic mouse models representing the main molecular and phenotypic subtypes of human melanomas and exhibiting their range of responses to immune checkpoint blockade (ICB). Comparative analysis of genomic, transcriptomic and tumor-infiltrating immune cell profiles demonstrated alignment with clinical observations and validated the correlation of T cell dysfunction and exclusion programs with resistance. Notably, genome-wide expression analysis uncovered a melanocytic plasticity signature predictive of patient outcome in response to ICB, suggesting that the multipotency and differentiation status of melanoma can determine ICB benefit. Our comparative preclinical platform recapitulates melanoma clinical behavior and can be employed to identify new mechanisms and treatment strategies to improve patient care.

---

Immune checkpoint blockade (ICB) has become the first-line treatment for metastatic melanoma, the deadliest skin cancer. Antibodies inhibiting CTLA-4 and PD-1/PD-L1 signaling pathways have been approved, as monotherapies or in combination, for more than ten cancer types in this decade due to their significant improvement of patient survival^1^. Yet, even in melanoma, the “poster child” for ICB success, response rates are insufficient and alternative strategies are being explored, including combinations with chemotherapy, radiotherapy, targeted therapy, and other immune checkpoint inhibitors^2^. Despite the intensive efforts to enhance ICB efficacy, understand mechanisms of sensitivity/resistance, and discover predictive biomarkers, the determinants of ICB response are still poorly understood. High mutation and neoantigen burden in pretreated tumors, increased T cell infiltrates and induction of inflammatory pathways during treatment, and antigen presentation alterations have shown correlation with clinical benefit^3–18^. Although the expression of specific pathways has been associated with ICB response in particular patient cohorts^7,18,19^, uncovering broad predictive signatures based on gene expression profiles has been elusive due to difficulties in collecting high quality transcriptomic data from clinical sets. Recently, independent computational predictors were developed based on immune-related gene expression profiles, such as immune checkpoints, co-stimulatory molecules and T cell dysfunction and exclusion markers^20–22^. However, the utility of these approaches for patient stratification will require further validation.

Mouse models have historically served as essential tools for plumbing mechanisms underlying tumor initiation, progression and drug response, and have enormous potential. However, their value for informing clinical trials and predicting patient outcome is still much debated, and likely depends on the quality and diversity of the models employed. Although wildtype mice rarely develop melanomas, they have been genetically engineered with oncogenic drivers to successfully provoke melanomagenesis^23–27^. Nevertheless, the creation of reliable melanoma models is handicapped by the inherent differences between mouse and human skin architecture and the complexity and heterogeneity of the human disease. Moreover, the majority of genetically engineered mouse (GEM) models fail to incorporate appropriate exposure to ultraviolet (UV) radiation, known to be the major etiological melanoma risk factor and speculated to enhance susceptibility to immunotherapy^28,29^, likely contributing to the low mutation burden and poor immunogenicity that characterize most models^30–32,33^. These differences likely compromise the relevance and validity of current preclinical mouse studies.

We have reported that constitutive activation of the receptor tyrosine kinase MET in hepatocyte growth factor transgenic mice (*HGF^tg^*) leads to the human-like localization of melanocytes within or near the epidermis, which are then susceptible to melanoma induction by a single burning dose of neonatal UV radiation^34,35^. Melanomas arising under these circumstances are highly reminiscent of human melanomas, and our base *HGF^tg^* model can be combined with a variety of other oncogenic drivers to replicate the molecular diversity of human melanoma^36^. Here we report the generation and characterization of a new panel of melanoma GEM models for the preclinical study of immunotherapies. The comparative analyses of the mutational landscapes, transcriptomes, and tumor-infiltrating immune cell profiles from these models, in concert with cross-validation using clinical datasets, demonstrated the reliability of the platform and uncovered a novel melanocytic plasticity signature (MPS) predictive of ICB efficacy in patients.

## RESULTS

### Modeling diverse subtypes of human melanoma in mice

Aiming to recapitulate melanoma diversity, we developed four syngeneic models in C57BL/6 mice harboring a variety of clinically relevant genetic modifications and exposed to different carcinogens (**Supplementary Fig. 1**). Melanoma model 1 (M1) was induced in *Braf^CA/+^*; *Pten^flox/+^*; *Cdkn2a^flox/+^*; *Tyr-Cre^ERT2-tg^* mice by UV radiation at postnatal day 3 and activation of Cre^ERT2^ at day 7^23,34,37^. Melanoma model 2 (M2) was generated in *Braf^CA/+^*; *Cdkn2a^flox/+^*; *Tyr-Cre^ERT2-tg^*; *Hgf^tg^* mice and induced as in M1. Melanoma model 3 (M3) was induced in *Cdk4^R24C/R24C^*; *Hgf^tg^* mice by topic administration of 7,12-Dimethylbenz(a)anthracene (DMBA) at day 3^38,39^. Finally, Melanoma model 4 (M4) was generated in *Hgf^tg^* mice by UV radiation at day 3^34^. Tumor fragments from each model were expanded in C57BL/6 mice as GEM-derived allografts (GDAs^40^) and viably cryopreserved to generate a tumor biobank. In parallel, cell lines from each model (CL1-4) were established and implanted into syngeneic mice (CLDA) for complementary studies (**Supplementary Fig. 1**).

We characterized the mutational landscape of the four models by whole exome sequencing (WES) analysis of the cell lines and GDAs. We found the highest number of non-synonymous single nucleotide variants in M1 and M4 and the lowest in M2 (**Fig. 1a; Supplementary Table 1**). We also performed spectral karyotyping (SKY) of the four cell lines to detect additional alterations such as chromosomal rearrangements or aneuploidy. This analysis showed 60% of tetraploid cells in M1 cell lines and a high number of translocations and chromosomal duplications in M3 and M4, whereas no obvious alterations were observed in M2 (**Supplementary Fig. 2a; Supplementary Table 2**). Consistent with the melanoma-inducing carcinogens C>T transitions, characteristic of COSMIC UV-related signature 7^41^, were predominant in M1, M2, and M4; in contrast, A>T transversions, characteristic of DMBA exposure, were the main mutations found in M3 (**Fig. 1b,c**). Notably, oncogenic driver and tumor suppressor mutations often detected in human melanomas^42,43^ (**Fig. 1d**, right panel) occurred spontaneously in the models, including *Kras* (M4), *Gnaq/z/11* (M2, M3, and M4), *Sf3b1* (M1 and M4), *Erbb4* (M1 and M4) and *Trp53* (M1 and M3) (**Fig. 1d**, left panel; **Supplementary Table 3**). We next clustered the four models together with the TCGA melanoma dataset according to their mutation profiles^36,43^. M1 and M2 clustered together with BRAF-mutant patients, M3 was in the proximity of triple-wildtype (*BRAF/RAS/NF1*-wildtype) and M4 clustered with RAS mutant melanomas (**Supplementary Fig. 2b**). We confirmed by western blot the activation of MAPK and PI3K pathways downstream of the genetic modifications incorporated in our models (**Supplementary Fig. 2c**). M3 and M4 exhibited overt MET phosphorylation, likely a consequence of the *Hgf* transgene activity. ERK phosphorylation was enhanced in *Braf^CA/+^*-harboring M1 and M2, followed by M4 (containing *Kras^G12D^* mutation). Marked activation of AKT by *Pten* heterozygous deletion in M1 was also noted. Altogether these results suggest that M1 and M2 represent subsets of BRAF mutant, M4 of RAS mutant and M3 of triple-wildtype patient melanomas.

**Fig. 1:**
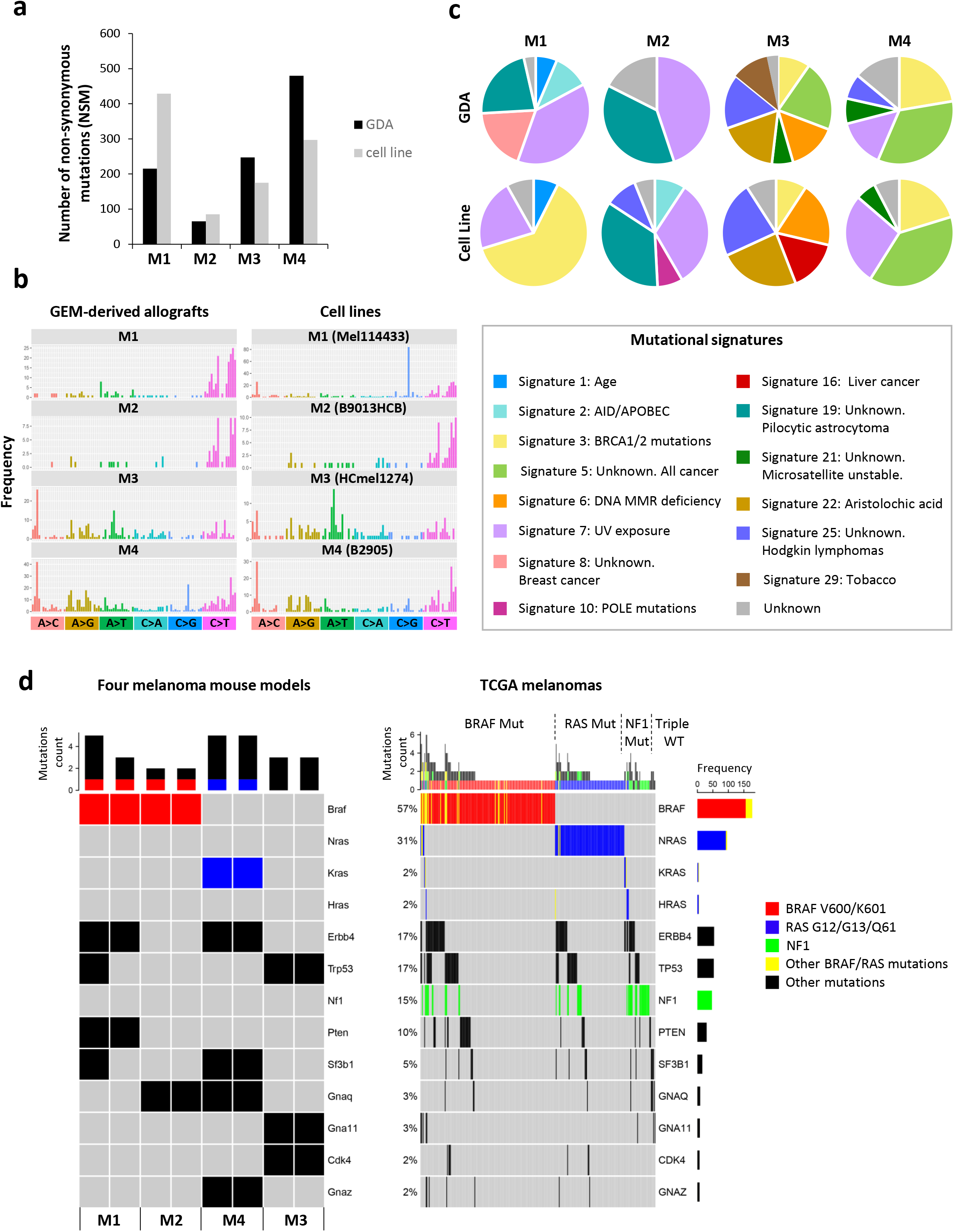
Modeling diverse subtypes of human melanoma in mice. **a**, Number of non-synonymous mutations (NSM) obtained from whole-exome sequencing of GEM-derived allografts (GDA, black bar) and cell lines (grey bar) from the four melanoma models (M1-M4). **b**, Frequency of the indicated single nucleotide mutations found in the four models (GDAs and cell lines). **c**, Distribution of COSMIC mutation signatures^41^ found in the four models (GDAs and cell lines). **d**, Hot spot mutations frequently found in human melanomas (right panel) that were detected in the four models (left panel). Each mouse model (M1-M4) depicts 2 columns, representing results from either the cell line (left) or GDA (right). BRAF, RAS and NF1 alterations are highlighted as they are used to classify the different molecular subtypes of human melanomas^36,43^.

### The melanoma mouse models recapitulate patient diversity in response to CTLA-4 blockade

To evaluate the response of our models to immune checkpoint blockade, we implanted GDAs or CLDAs from each model into C57BL/6 mice and treated them with either anti-CTLA-4 (αCTLA-4) or isotype control antibodies (**Fig. 2a**). αCTLA-4 treatment did not affect M1 and M2 tumor growth nor survival. Conversely, 20-30% of M3 and 40% of M4 melanomas treated with αCTLA-4 reproducibly showed tumor growth delay resulting in significantly improved survival for M4-bearing mice (**Fig. 2b-e**). Overall, our models displayed a broad range of responses to αCTLA-4 from resistant M1 and M2 to partially and highly sensitive M3 and M4, respectively, mimicking the diverse clinical responses found in human melanoma.

**Fig. 2:**
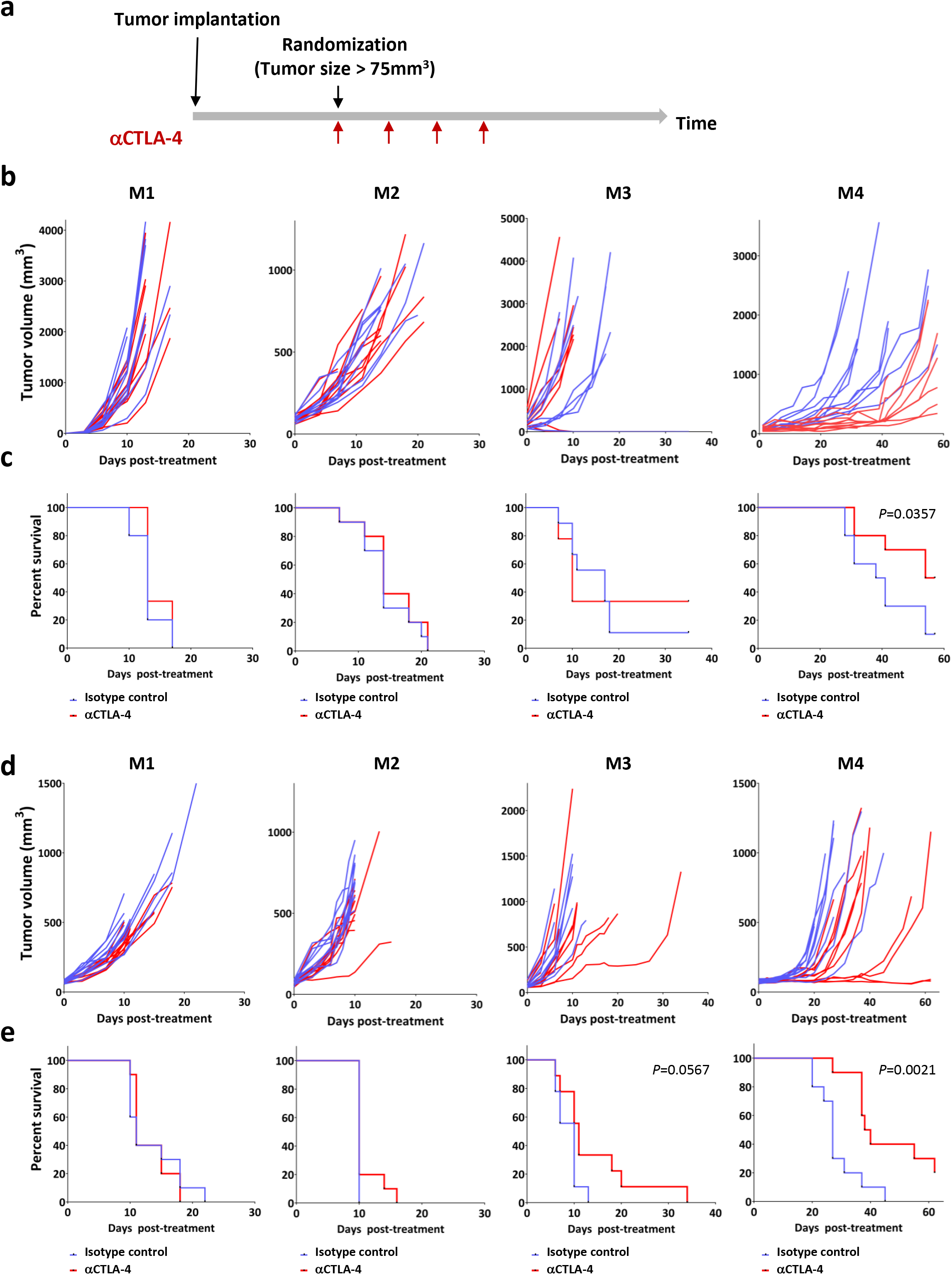
The melanoma mouse models recapitulate patient diversity in response to CTLA-4 blockade. **a**, 2-3 mm^3^ tumor pieces (GDA) or 1.0 × 10^6^ melanoma cells (cell line) from each model were subcutaneously (s.c.) implanted into C57BL/6 syngeneic mice. When tumors reached 75-125 mm^3^, mice were randomized and treated with anti-CTLA-4 (aCTLA-4) or isotype antibody as indicated. **b**, Tumor growth of the four melanoma models (GDAs) upon treatment with aCTLA-4 (red lines, N=10) or isotype control (blue lines, N=10). **c**, Kaplan-Meier survival curves from (b). **d**, Growth curves of the four melanoma models (cell line-derived allografts) after treatment with aCTLA-4 (N=10) or isotype control (N=10). **e**, Kaplan-Meier survival curves from (d). *P*-values from Log-rank (Mantel-Cox) test are indicated (c and e).

ICB efficacy depends on the anti-tumor immune responses generated upon recognition of the cancer cells^1^. To determine the immunogenicity of the melanomas we performed vaccination assays *in vivo*^44^. γ-Irradiated cells from each model were subcutaneously injected into C57BL/6 mice. Four weeks post vaccination, untreated cells from the same model were implanted into the opposite flank of the animals (**Supplementary Fig. 3a**). Vaccination had no significant effect on the tumor-free survival of the mice challenged with either M1 or M2 cells when compared with non-vaccinated control mice. In contrast, vaccination significantly delayed tumor onset in the mice challenged with M3 or M4 cells (**Supplementary Fig. 3b-e**). These results suggested that tumor immunogenicity correlates with the response to αCTLA-4.

### Antigen presentation pathway is functional in the four melanoma models

Altered antigen presentation in tumor cells has been proposed as a mechanism of clinical resistance to ICB^12,45^. Notably, no mutations in antigen presentation genes were found in any of our melanoma models (**Supplementary Table 1**). In addition, we confirmed the induction of major histocompatibility complex (MHC) class I-associated genes following IFNγ stimulation in all cell lines, and we validated the expression of H2-Kb protein by flow cytometry (**Supplementary Fig. 4a,b**). Furthermore, when we analyzed the transcriptome of the four melanomas either untreated or after CTLA-4 blockade, no significant differences were found in the expression of MHC class I- and class II-related genes (**Fig.3a; Supplementary Table 5**). This suggests that the integrity of the antigen presentation pathway is maintained in all the models.

**Fig. 3:**
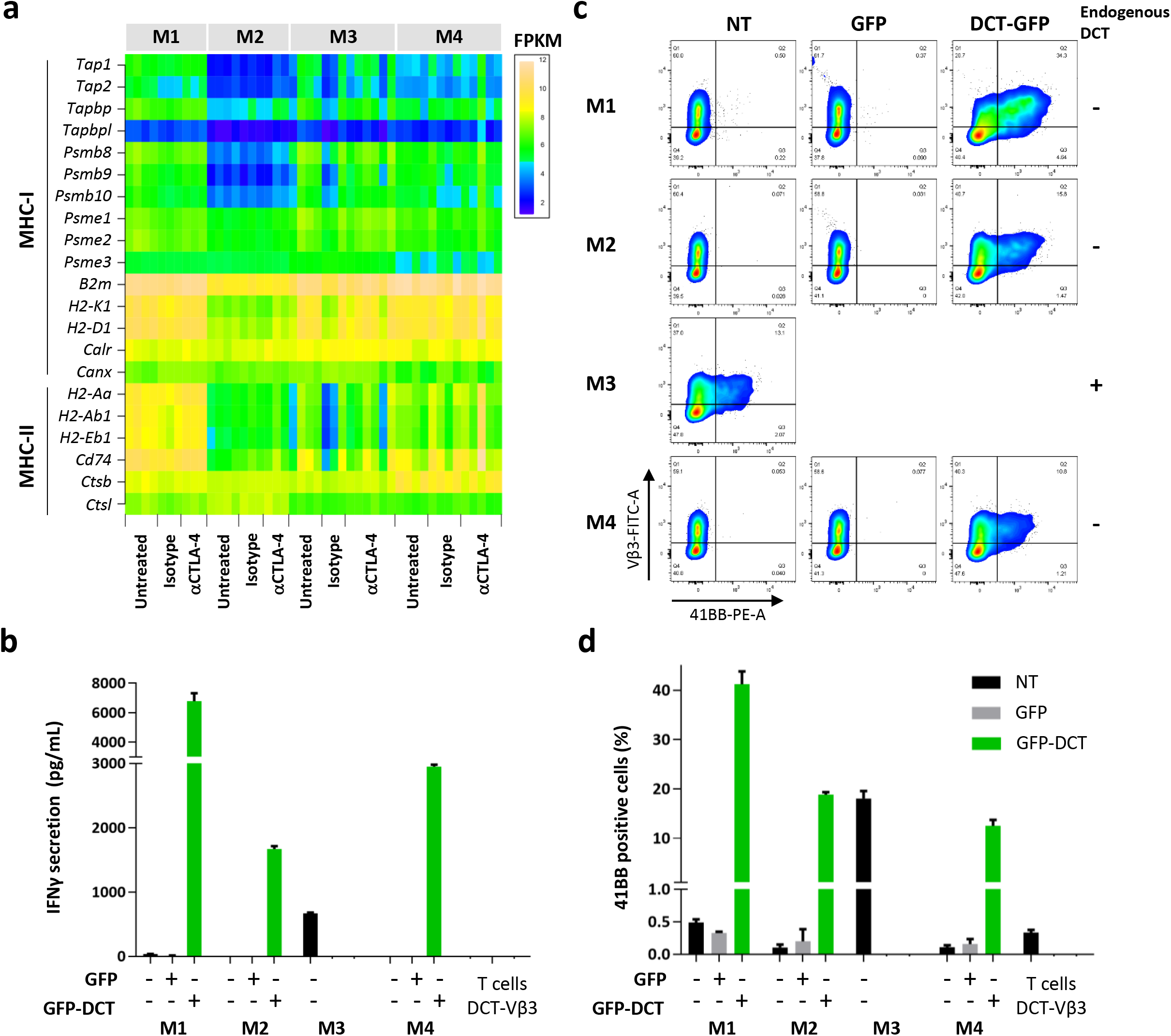
Antigen presentation is functional in the four melanoma models. **a**, Expression of the indicated major histocompatibility complex (MHC) class I and class II related genes in the four models. The heatmap depicts the FPKM obtained from RNA sequencing of untreated (N=4), isotype control (N=3 for M1-3, N=4 for M4) or aCTLA-4 treated (N=3 for M1 and M2, N=6 for M3 and N=5 for M4) cell line-derived allografts. **b**, IFNγ concentration in the media from 24-hour co-cultures of DCT-reactive T cells and the four models cells expressing GFP-DCT peptide ^46,47^, GFP empty vector or non-transduced. CD8^+^ sorted T cells from C57BL/6 splenocytes were transduced with recombinant TCR Vβ3 for the recognition of 9 aa DCT peptide (DCT-Vβ3) or Thy1.1 as control. Graph depicts IFNγ pg/mL measured by ELISA. **c**, T cell activation after 24-hour co-culture with the indicated melanoma cell lines measured by 41BB expression analyzed by flow cytometry. **d**, Percentage of 41BB positive cells obtained from (c). Co-cultures were done in triplicates and data is shown from a representative of 2 independent experiments.

To further evaluate the functionality of the pathway, we co-cultured cells from each model with engineered CD8^+^ T cells reactive against a specific antigen from the melanocytic marker dopachrome tautomerase (DCT). Only M3 cells expressed substantial levels of *Dct* in culture conditions; therefore, we overexpressed the 9aa DCT peptide^46^ recognized by the engineered T cells in M1, M2 and M4 cells (**Supplementary Fig. 4c**). The 9aa DCT peptide was fused to the C-terminus of eGFP protein but separated by a 2aa linker to ensure proper cleavage and presentation by MHC-I^47^. We used ELISA to measure IFNγ concentration in the media of 24-hour co-cultures as readout of T cell activation. IFNγ was accumulated 100- to 1000-fold in the media of DCT-expressing cells from all models when co-cultured with DCT-reactive T cells, but not in the presence of Thy1.1-transduced control T cells (**Fig. 3b; Supplementary Fig. 4d**). Moreover, 15-40% of the DCT-reactive T cells, but not Thy1.1 control, were activated by DCT-expressing cells from the four models as evidenced by the expression of the activation marker 4-1BB (CD137) measured by flow cytometry (**Fig. 3c,d; Supplementary Fig. 4e**). These results confirmed the intact ability of the melanoma cells from the four models to present antigens in the context of MHC-I and activate CD8^+^ T cells.

### Correlation of tumor-infiltrating lymphocytes with αCTLA-4 response in the melanoma models

Increased T cell infiltration early on or after treatment has been reported to correlate with ICB efficacy, although baseline tumor infiltrating lymphocytes (TILs) do not predict response^5,7,48,49^. We first examined TILs in the four models by CD3 immunostaining of untreated melanomas (**Supplementary Fig. 5a**). Automated quantification of the total CD3 positive area per section did not reveal a clear distinction between sensitive and non-responsive models, with resistant M1 and sensitive M4 displaying the highest T cell infiltration (**Supplementary Fig. 5b**). Next, we addressed T cell infiltration after αCTLA-4 treatment of M4. We harvest the tumors at early time points when progressively growing could be distinguished from stable/regressing melanomas, but size was comparable (21-28 days, **Supplementary Fig. 5c**). Consistent with previous clinical observations, those tumors that responded to αCTLA-4 exhibited significantly increased TIL densities relative to those not responding or treated with isotype control (**Supplementary Fig. 5d**).

We next performed in depth characterization of the immune cell populations infiltrating untreated melanomas from our models by unbiased high parametric flow cytometry analysis. Although we only observed a modest decrease of the total number of infiltrating leukocytes (CD45^+^ cells) in M4, we found a strikingly distinct distribution of the immune populations between the four models (**Fig 4a-c**). The lymphoid fraction was enriched in M4, followed by M1, while macrophages and neutrophils were predominant in M2 and M1 (**Fig 4c; Supplementary Fig. 6a-d**). In contrast, dendritic cells (DCs) were more abundant in M3 (**Supplementary Fig. 6e**). In agreement with the immunostaining results, CD3^+^ T cells were highest in M4 and M1, almost absent in M2 (≤ 3%) and intermediate in M3 (**Fig. 4d**). CD8^+^ T cells represented the majority of the T cell compartment in M1 and were found in similar frequencies in M4 (**Fig. 4e**). Total CD4^+^ T cells were more abundant in M4 and M3, and although the frequency of regulatory T cells (CD3^+^CD4^+^CD25^+^FoxP3^+^, T reg) was also higher in the same tumors, they represented a small proportion of total CD4^+^ cells (**Fig. 4f; Supplementary Fig. 6f**). Notably, the scarce T cells found in the resistant M2 tumor were predominantly T reg (**Supplementary Fig. 6f**). Consistent with their higher CD8^+^ T cell content, CD8^+^ T cell/T reg ratios were increased in M1 and M4 (**Fig. 4g**). Of note, unconventional γδ T cells, which exert anti-tumor cytotoxic responses^50^, were rare but higher in the sensitive M4 (**Fig. 4h**). Altogether, these results revealed the immunological diversity of the models and suggested that in addition to quantity, the quality of TILs and the presence of other immune cell populations may be an important determinant of response.

**Fig. 4:**
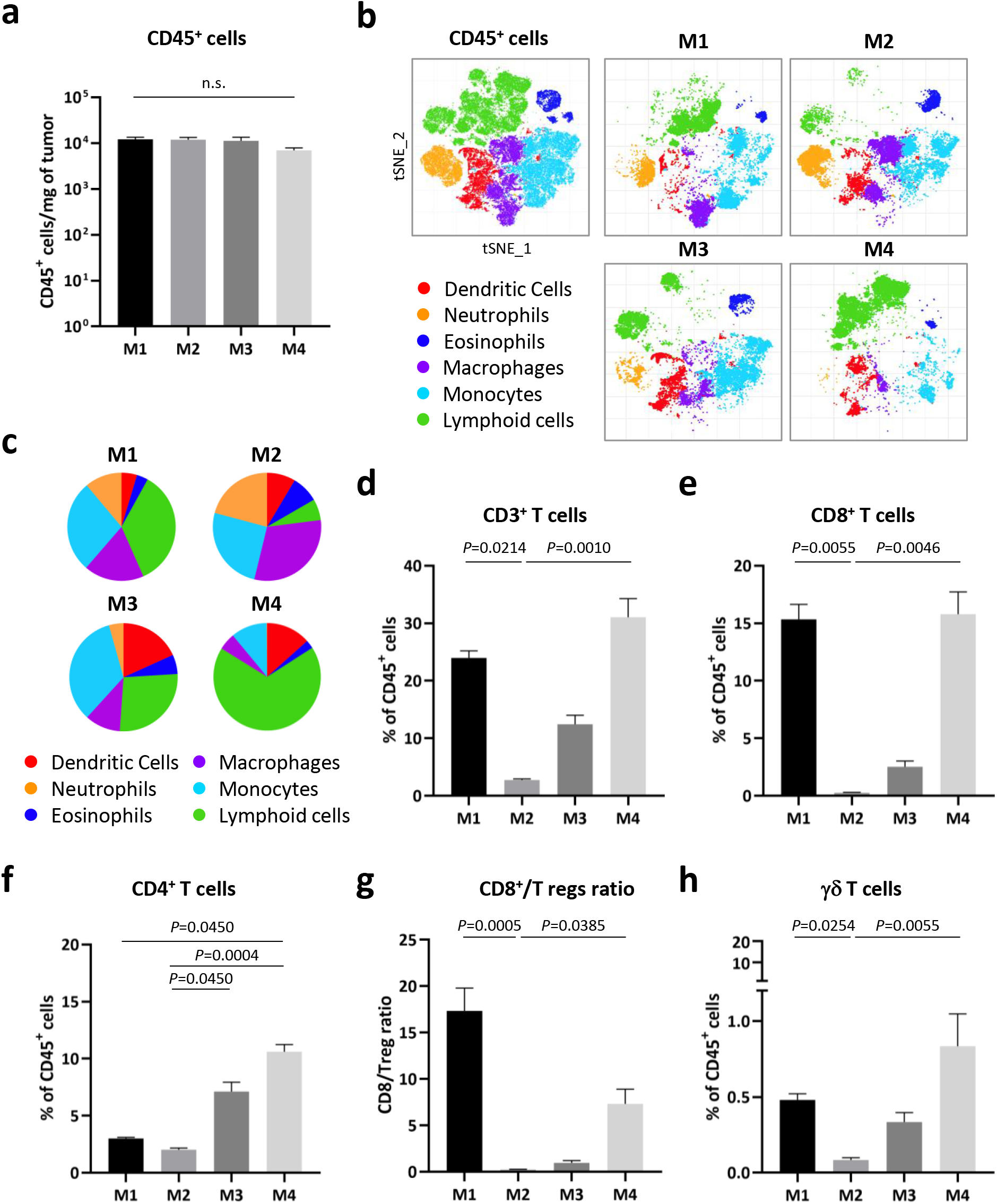
Intratumoral immune cell correlates of aCTLA-4 response in the melanoma models. **a**, Total number of leukocytes (CD45^+^ cells) infiltrating the untreated cell line-derived allografts of the four melanoma models measured by flow cytometry. **b**, 20-parameter flow cytometry analysis by t-stochastic neighbor embedding (t-SNE) of the intratumoral CD45^+^ cells from (a). **c**, Pie charts showing the indicated immune cells fractions obtained from (b). **d-f**, Percentage of the intratumoral CD3^+^ (d), CD8^+^ (e), CD4^+^ (f) in the four models (untreated). **g**, Ratio of intratumoral CD8^+^ cells/Tregs (CD3^+^TCRβ^+^CD4^+^CD25^+^FoxP3^+^) from (a). **f**, Percentage of γδ T cells from (a). Data from a representative of two experiments is depicted as the mean (N=5) and error bars represent S.E.M. Kruskal-Wallis test *P*-values adjusted for multiple comparisons are indicated (a,d-h). n.s.: non-significant. See also **Supplementary Table 4** for the gating description.

### The models resistant to αCTLA-4 exhibit T cell dysfunction and exclusion profiles

Overall the frequency and number of TILs were not enough to explain the different response to αCTLA-4 of resistant M1 and sensitive M4. To further address the mechanism of immune evasion we analyzed the expression of T cell exhaustion markers^51,52^ from the RNA sequencing data of the four models. The expression levels of the markers in M1 highly correlated with T cell exhaustion profiles (**Fig. 5a**). Next, we used t-stochastic neighbor embedding (t-SNE) analysis of the high parametric flow cytometry data to decipher the CD3^+^ T cell populations infiltrating the four models and confirmed that M1 had a higher proportion of CD8^+^ exhausted T cells (**Fig. 5b,c; Supplementary Fig. 7a**). Moreover, the level of PD-1, LAG3 and TIM-3 expression was also significantly elevated in M1 CD8^+^ T cell infiltrates (**Supplementary Fig. 7b-d**). These data evidenced a predominant T cell exhaustion phenotype in M1, which may explain the high TMB and TIL densities but resistance to αCTLA-4 exhibited by this model.

**Fig. 5:**
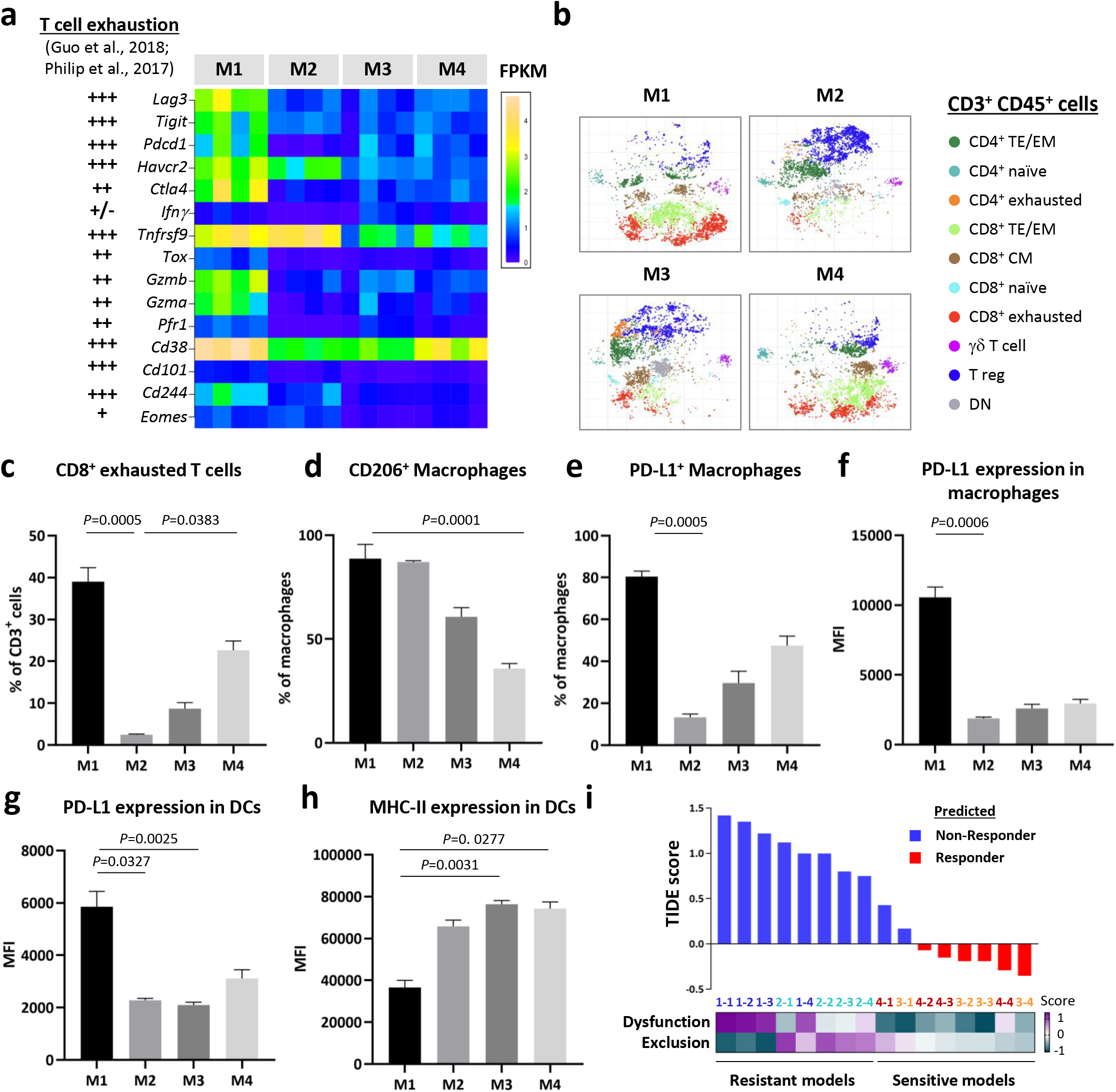
The models resistant to aCTLA-4 exhibit T cell dysfunction and exclusion profiles. **a**, Expression of T cell exhaustion and dysfunction markers ^52,53^ in the cell line-derived allografts from the four models. Data is shown as FPKM from RNA sequencing analysis (untreated, N=4). **b**, t-stochastic neighbor embedding (t-SNE) analysis of the intratumoral CD3^+^ lymphocytes in the four models (untreated, N=5). TE/EM: T Effector/Effector memory (CD62L^−^ CD44^+^); CM: central memory (CD62L^−^ CD44^+^); naïve (CD62L^+^CD44^−^); exhausted (PD-1^+^ LAG3^+^TIGIT^+^TIM3^+^) and DN: double negative (CD4^−^CD8^−^). **c-e**, Percentage of intratumoral exhausted CD8^+^ T cells (c), CD206^+^ (d) and PD-L1^+^ (e) macrophages (CD68^hi^CD64^+^;F480^+^Ly6C^−^) in the four models. **f-h**, Expression of PD-L1 (f,g) and MHC-II (h) in macrophages and dendritic cells (DCs, CD11c^+^ MHC-II^+^ CD64^−^ CD24^int/hi^ CD135^+^) as the mean fluorescence intensity (MFI) from flow cytometry. Data from a representative of two experiments is depicted as the mean (N=5) and error bars represent S.E.M. Kruskal-Wallis test *P*-values adjusted for multiple comparisons are indicated (c-h). See also **Supplementary Table 4** for the gating description. **g**, Tumor Immune Dysfunction and Exclusion (TIDE) scores^21^ of the four models (untreated, N=4). Blue bars represent the melanomas predicted as “Non-responders” and red bars those as “Responders”. Each tumor is color coded as: M1=dark blue, M2=light blue, M3=orange and M4=red.

We also characterized the myeloid compartment in the tumor microenvironment (TME) of the models. Most of the macrophages found in M1 and M2 were CD206^+^, a marker widely used to identify tumor-promoting macrophages (**Fig. 5d**). Moreover, M1 was also enriched in macrophages expressing the highest levels of PD-L1 (**Fig. 5e,f**). DCs in M1 tumors in addition to being at a lower frequency (<2% vs. 6–9% in M4; **Supplementary Fig. 6f**), expressed higher PD-L1 and reduced MHC-II levels (**Fig. 5g,h**). These findings suggest that the myeloid cell compartment supports the T cell dysfunction observed in M1 melanomas. Of note, the intratumoral macrophages of M2 expressed significantly high CD206 and RNA sequencing analysis by CIBERSORT^53,54^ confirmed a trend toward enrichment of the pro-tumor macrophage fraction in M1 and M2 (**Supplementary Fig. 8a,b**). Additionally, the expression levels and fraction of MHC-II^+^ macrophages were significantly decreased in M2, which correlated with higher β-Catenin mRNA expression in this model (**Supplementary Fig. 8c,d**). That could contribute to an impaired T cell recruitment/expansion into M2 melanomas, as described before in preclinical and clinical studies^16,55^. Overall, these results revealed a distinct immune suppressive TME in the αCTLA-4-resistant M1 and M2 models.

T cell dysfunction and exclusion signatures were recently implemented in a computational method (Tumor Immune Dysfunction and Exclusion, TIDE) that could predict αCTLA-4 and αPD-1 response in melanoma patients from three independent data sets^21^. Importantly, by applying this tool to the RNA expression data from our models, 100% of the untreated M1 and M2 were predicted as “non-responders” and 75% of M3 and M4 as “responders” by a cutoff value of 0 (**Fig. 5i**). Moreover, M1 exhibited high dysfunction scores while M2 had increased exclusion scores. These results support the key role of T cell dysfunction and exclusion programs in the resistance to ICB and highlight the utility of our models for validating computational predictive methods.

### Transcriptomic profiling of the models identifies a melanocytic plasticity signature that predicts patient outcome in response to ICB

To explore the gene expression profiles that may influence the response to ICB we analyzed RNA sequencing data from the four untreated melanoma models by pairwise differential gene expression and Gene Set Enrichment Analysis (GSEA) (**Supplementary Tables 5, 6**). We observed an upregulation of “melanocytic markers” in M3 and M4, whereas “nervous system-related”, “Epithelial-Mesenchymal Transition (EMT)” and “inflammation” genes were prevalent in M1 and M2 (**Supplementary Fig. 9a**). We used immunostaining of untreated mouse melanomas and we validated the expression of the melanocytic markers DCT and tyrosinase related protein 1 (TYRP1) only in M3 and M4, and expression of the neural crest lineage transcription factor SOX10 in all the models, indicative of a distinct differentiation status of the models (**Supplementary Fig. 9b**).

Next, we developed a gene signature predictive of ICB efficacy through a comparison of the resistant M1 and M2 versus the sensitive M3 and M4 transcriptomes. Based on differential gene expression statistics and sequential evaluation of patient response prediction by Fisher’s exact test using a metastatic melanoma cohort treated with αCTLA-4 (Van Allen data set^4^), we identified a 45-gene signature comprised of 33 upregulated and 12 downregulated genes in M1/M2 (see **Methods; Supplementary Table 7**). To examine the biological functions of the genes in the signature we performed Ingenuity Pathway Analysis (IPA) and found “nervous system development and function” and “neurological disease” as the top 2 most significant categories (**Supplementary Fig. 10a**). Since melanocytes and peripheral nervous system have a common embryonic origin in the neural crest and share stem cell progenitors in the skin^56^, we next explore the signature expression in several melanocytic precursors. We computed scores based on the expression of the 45 genes in embryonic mouse melanoblasts at day E15.5 and E17.5 and differentiated melanocytes from pups at postnatal day 1 and 7 (P1 and P7, respectively; Marie et al., BIORXIV/2019/721712). The signature expression in melanoblasts was aligned with resistant M1/M2 while melanocytes aligned with sensitive M3/M4 (**Supplementary Fig. 10b**). We also interrogated the signature expression in two hair follicle melanocytic stem cell populations from the adult mouse skin (P56) with distinct regenerative capacities^57^. Consistently, the multipotent progenitors with neural crest stem cell-like properties (CD34^+^) showed higher scores than those committed to melanocytic differentiation (CD34^−^, **Supplementary Fig. 10c**). Notably, CD34 expression was high in resistant M1/M2 and low in sensitive M3/M4 (data not shown). Altogether, these results suggest that the signature reflects the potency and differentiation status of the melanocytic lineage, which we will hereafter refer to as a “Melanocytic Plasticity Signature” (MPS). High MPS scores represent undifferentiation, multipotency and/or neural crest stem cell properties and low MPS scores, later stages of melanocytic differentiation.

We next analyzed MPS expression in the Van Allen data set^4^ and found significantly lower scores in responders compared to non-responders (**Fig. 6a**). Importantly, MPS scores correctly predicted the response of 81% of the patients using 33 percentile as a cutoff (Fisher’s exact *P*-value=0.00489, **Fig. 6b**). Furthermore, high MPS scores significantly correlated with worst progression-free survival (PFS) and overall survival (OS) of the patients in this cohort (**Fig. 6c,d**). To validate our results in an independent ICB data set we compiled baseline RNA sequencing data of immunotherapy naïve patients from two metastatic melanoma cohorts treated with αPD-1 (Hugo and Riaz data sets^8,19^). In concert with the Van Allen data set, non-responder melanomas exhibited significantly higher MPS scores than responders (**Supplementary Fig. 11a**). Moreover, low-MPS score patients showed significantly better OS (**Supplementary Fig. 11b**). These results demonstrate the value of the MPS for predicting patient outcome in response to ICB and suggest that the potency and differentiation status of the melanomas could determine ICB efficacy.

**Fig. 6:**
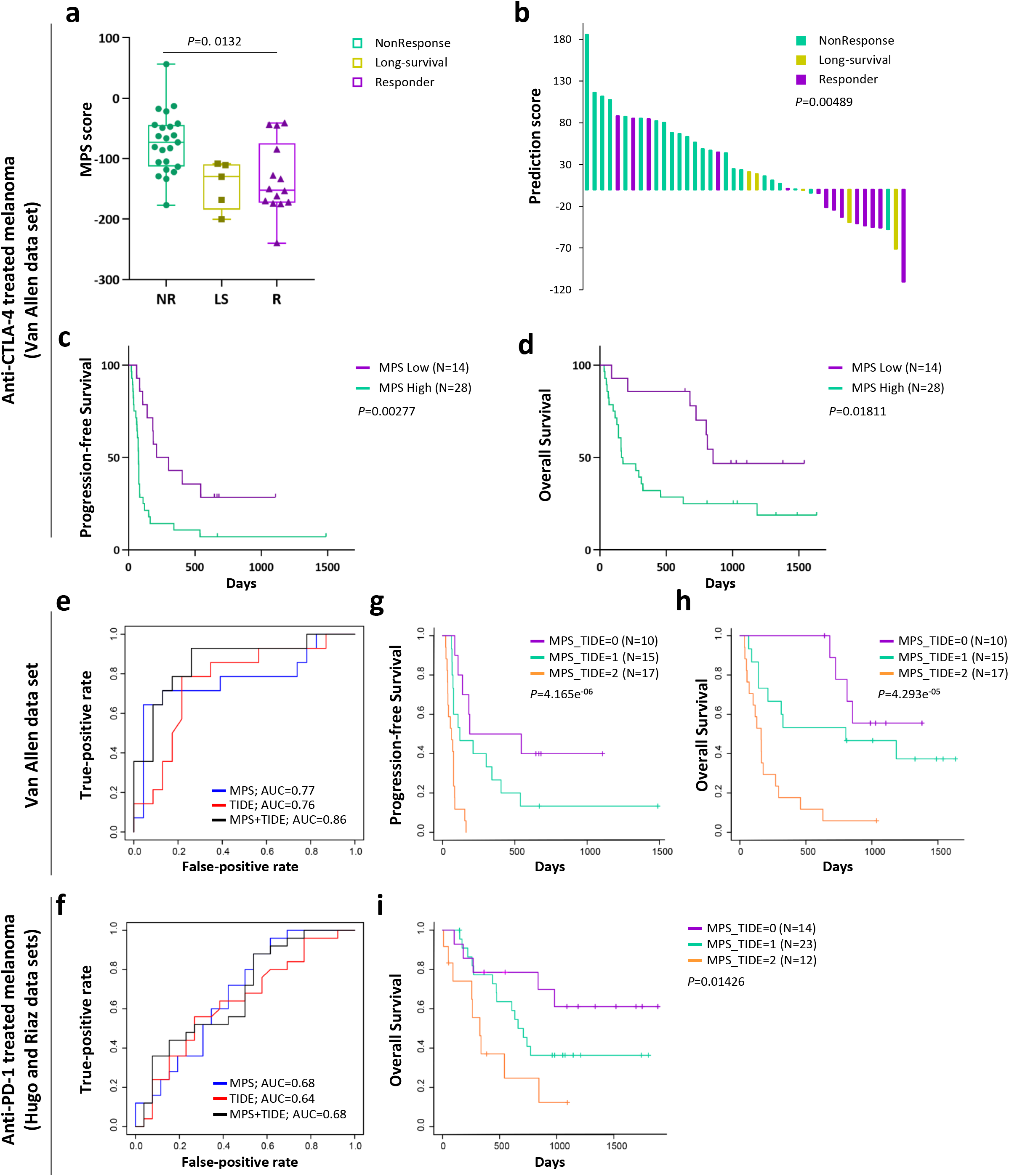
Transcriptomic profiling of the models identifies a melanocytic plasticity signature that predicts patient outcome in response to ICB. **a**, Expression of the melanocytic plasticity signature (MPS) in metastatic melanoma patients treated with ipilimumab (aCTLA-4, Van Allen data set^4^). The box plot shows the MPS scores in pretreated melanomas from non-responder (NR, green, N=23), long-survivors but non-responders by RECIST (LS, yellow, N=5) and responder (R, violet, N=14) patients. Kruskal-Wallis test *P*-value adjusted for multiple comparisons is indicated. **b**, Waterfall plot representing prediction scores for each patient in Van Allen data set. Prediction significance was analyzed by Fisher’s exact test and *P*-value is indicated. **c-d**, Kaplan-Meier curves of the progression-free (c) and overall (d) survival for ipilimumab-treated patients accordingly to their MPS scores. **e-f**, ROC curves to evaluate the prediction performance of MPS, TIDE and MPS+TIDE in Van Allen (e) and Hugo-Riaz (f) data sets. The area under the ROC curves (AUC) values for each predictor are indicated. **g-i**, Kaplan-Meier curves of the progression-free (g) and overall (h,i) survival for Van Allen (g,h) and Hugo-Riaz (i) data sets by the combination of MPS and TIDE scores. MPS_TIDE=0: MPS^low^ TIDE^low^; MPS_TIDE=1: MPS^high^ TIDE^low^ or MPS^low^ TIDE^high^; and MPS_TIDE=2: MPS^high^ TIDE^high^. N for each group is indicated and *P*-values calculated by the Log-rank (Mantel-Cox) test (c-d and g-i).

### The combination of MPS and TIDE scores improves the prediction of patient response and survival upon ICB in multiple data sets

To address whether the predictive power of MPS was associated with reported immune biomarkers we analyzed the expression of CTLA-4, PD-1, PD-L1, PD-L2 and the cytolytic activity index in the Van Allen data set^4^ and found no difference between high- and low-MPS score patients (data not shown). Next, we employed receiver operating characteristic (ROC) curves to compare the prediction performance of MPS with PD-L1 expression and TMB, used as biomarkers of αPD-1 response for some cancers^58^, and the TIDE method. Importantly, MPS and TIDE achieved the best AUC values in the Van Allen and Hugo-Riaz data sets (**Supplementary Fig. 11c**). We hypothesized that the combination of MPS and TIDE scores would further improve the ability to predict ICB efficacy. Because both predictors have different scale ranges, we calculated z-scores for each one and aggregated them to obtain combined MPS+TIDE scores (see **Methods**). ROC curves showed a marked improvement of the AUC values by MPS+TIDE for Van Allen cohort but the same as MPS alone for Hugo-Riaz data set (**Fig. 6e,f**).

To evaluate the effect of each signature on survival we used Cox proportional hazards model (Cox-PH) analysis. MPS and TIDE showed significant hazard ratios (HR) for the PFS and OS of the Van Allen data set as well as for the OS of the Hugo-Riaz data set (**Supplementary Table 8**). Notably, the significance was maintained in multivariate Cox-PH analysis containing both MPS and TIDE, suggesting they can have an additive effect when combined (**Supplementary Table 9**). We next stratified the melanomas from each data set into three groups accordingly to their MPS and TIDE scores and found significantly improved PFS and OS for low-MPS and low-TIDE group (MPS_TIDE=0, violet), whereas patients with high scores for both predictors (MPS_TIDE=2, orange) exhibited the poorest survival in the Kaplan-Meier curves (**Fig. 6g-i**). These results confirmed a better correlation with survival of MPS and TIDE combined versus each signature alone (**Fig. 6c,d; Supplementary Fig. 11b,d**). Our results demonstrate the power of combining cancer cell-intrinsic (potency and differentiation) and extrinsic (immune related) factors to build stronger predictors of patient survival in response to αCTLA-4 or αPD-1.

## DISCUSSION

In this study we characterized a set of four immunocompetent melanoma mouse models that faithfully represent the main molecular subtypes of human melanoma and recapitulate the diversity of the clinical response to ICB. In contrast to most preclinical studies based on single models, we utilized and compared multiple relevant GEM-derived melanomas that we cross-validated with human data sets, which allowed us to discern key factors predicting and/or influencing response and resistance. Analysis of tumor mutation burden, TILs densities and immune cell profiles demonstrated that our models resemble clinical associations with ICB response. Moreover, this new preclinical platform has provided insight into both melanoma cell extrinsic and intrinsic determinants of ICB efficacy. We identified T cell dysfunction and exclusion programs, as well as melanoma multipotency and differentiation status associated with resistance. Furthermore, generation and computational analysis of rich genomic and transcriptomic murine data sets uncovered a novel melanocytic plasticity signature (MPS) predictive of patient outcome in response to ICB and has provided a platform for the discovery of new biomarkers.

Our studies revealed a distinct immune composition in the TME of the four melanoma models, underlining non-overlapped immunosuppressive populations in resistant M1 and M2 models. We found, as in clinical observations, that TIL densities after CTLA-4 blockade correlated better with efficacy than at baseline. However, our results underscored the importance of the functional status of T cells over their number to distinguish responder from resistant tumors. The αCTLA-4 resistance of M1 despite high TMB and TIL quantities was explained by increased T cell dysfunction markers and elevated PD-L1 expression in its intratumoral macrophages and DCs. On the other hand, the enrichment of tumor-promoting macrophages, decreased MHC-II levels in DCs, and high expression of β-Catenin supported the “cold” TME of resistant M2. Moreover, we applied the recent TIDE method, which evaluates T cell dysfunction and exclusion profiles^21^, to untreated melanomas from our set of mouse models and obtained consistent predictions of their response, suggesting that they represent a suitable platform for the functional validation of predictions from patient data.

Our findings demonstrate that the application of mouse gene expression profiles to build predictors of ICB efficacy in the clinic, although challenging, is a valuable strategy for the discovery of evolutionarily conserved signatures. A clear advantage of the comparative analysis of distinct but genetically controlled syngeneic models is to reduce the noise derived from genetically diverse backgrounds, especially prevalent in human data sets. In contrast to current computational methods based on associations with bona fide immune-related genes^20,21^, we used genome-wide expression profiles to identify a signature representative of melanocytic plasticity, not previously linked to ICB or immune responses, that was predictive of ICB efficacy. Remarkably, the MPS predictor that was built to distinguish resistant M1/M2 from sensitive M3/M4 melanomas, also aligns with expression data from the Hornyak lab that distinguish two melanocyte stem cell (MSC) populations in the murine hair follicle with distinct multipotency capacities^57^. CD34^+^ neural crest-like MSCs correlated with resistant M1/M2, while a CD34^−^ MSCs that represent a more advanced state of melanocyte differentiation, corresponded to sensitive M3/M4. Melanoma differentiation could influence ICB efficacy by increasing the expression of tumor-self antigens that would potentiate immunogenicity, a phenotype clearly linked to response in our models. On the other hand, melanomas with a higher degree of stemness may in fact be more resistant to immunotherapy, as has been reported for resistance to chemo- and targeted-therapies^59,60^. Whether the MPS reflects melanoma immunogenicity, a melanoma stemness indicator, genes with an active immunomodulatory function, or a more complex combination of these remains to be elucidated in future studies. Regardless, our results underscore the complexity of ICB responses as the combination of immune-related (TIDE) and tumor cell intrinsic (MPS) factors demonstrated improved performance of patient outcome prediction.

We acknowledge that the validation of our signatures, although significant, is limited by the relatively small patient cohorts currently available that include pre-treatment samples from ICB naïve melanomas. This in fact emphasizes the importance of collecting high quality RNA sequencing data from the clinic and complementing them with preclinical data sets to develop accurate predictive methods. We anticipate that this new preclinical platform will prove useful for the identification of additional biomarkers and resistance mechanisms in the PD-1/PD-L1 pathway, for assessing the most efficacious adjuvant and neo-adjuvant regimens for ICB and for the discovery of new immunotherapeutic targets. Our studies demonstrate that mice can be engineered to be relevant and predictive for the treatment of human melanoma.

## METHODS

### Mouse models

All four mouse models in this study were generated on the C57BL/6 strain background, with the source of alleles described in the text and the associated references (**Table S1**). About 25% of Hgf^tg^ females develop prolapse at the age 6-8 months and have difficulties with pregnancies. Therefore, Hgf^tg^ females were not used for breeding. Both female and male mice were used for tumor induction. Cre^ERT2^ was activated by topical 4-hydroxy-tamoxifen (4-OHT) treatment as following: 0.1 ml of a 25 mg/ml solution in DMSO was administrated once daily for between 3 to 5 consecutive days starting at postnatal day 7 to a 1.5 × 1.5 cm patch of shaved dorsal skin ^61^. For UV treatment, a single dose of erythemal radiation generated from a bank of FS40 lamps was given to neonatal/perinatal transgenic mice (postnatal day 3 in this study) ^34^. M1, M2 and M3 melanomas were obtained from females and M4 melanoma from a male. Fragments from each melanoma model were expanded in syngeneic mice and viably cryopreserved to generate a GEM-derived allograft (GDA) biobank. The melanoma model 3 tumor (GDA3) and cell line (CL3) were kindly provided by Dr. Thomas Tueting laboratory (University of Magdeburg, Magdeburg, Germany) ^38,39^. GDA4 was generated by subcutaneous (s.c.) implantation of CL4 (B2905) and expansion in C57BL/6 mice for at least 3 passages.

### Cell lines

The cell lines from M1 (Mel114433), M3 (HCmel1274) and M4 (B2905) were cultured in RPMI supplemented with 10% of FBS and 1% L-Glutamine. The M2 cell line (9013BL) was cultured in Tu2% media (https://wistar.org/sites/default/files/2017-11/Herlyn%20Lab%20-%20Cell%20Culture%20Techniques%20-%202017.pdf). Authentication of all cell lines was performed by whole exome sequencing, genotyping the alleles in the corresponding mouse models, and Spectral karyotyping (SKY). CD8^+^ T cells from C57BL/6 spleens were cultured in RMPI supplemented with 10% FBS, 1% P/S, 50μg/mL Gentamicin, 1% L-Glutamine, 1% Sodium Pyruvate, 1% NEAA and 55μM β-mercaptoethanol at 37°C and 5% CO2. Cell lines were confirmed mycoplasma negative using Mycoplasma detection kit (Lonza, LT07-418).

### *In vivo* treatments and vaccination assays

All the mice used in the preclinical studies were 6-12 weeks old female C57BL/6N strain, supplied by Charles River facility in NCI-Frederick, with no selection on the weight. They were housed in AAALAC-accredited cage as a group of five, with *ad libitum* food and water, and 12-hour light cycle. When GDAs were used, a cryopreserved tumor fragment of the size 300-500 mm^3^ was divided and subcutaneously implanted into five mice. When the tumors reached an average size of 500 mm^3^, a tumor with exponential growth kinetics was selected to be divided and implanted into 10 mice. When these tumors reached an average size of 500 mm^3^, those with similar exponential growth kinetics were harvested, divided evenly, and transplanted into the mice planned for study.

For αCTLA-4 efficacy studies, 2-3 mm^3^ tumor pieces (GDA) or 1.0×10^6^ melanoma cells (CLDA) from each model were s.c. implanted into C57BL/6 mice. When the tumors reached 75-125 mm^3^, mice were randomized and treated with αCTLA-4 (BioXCell, BP0164) or isotype control (BioXcell, BP0086). Antibodies were administered intravenously (i.v.) at a final dose of 10mg per Kg twice per week for a total of 4 doses.

For vaccination assays, 1.0×10^6^ cells from each model received γ-irradiation at 35 (M1, M3 and M4) or 200 (M2) Gy using a ^137^Cs MARK I irradiator (JL Shepherd & Associates, San Fernando, CA), and then s.c. injected into dorsal surface of C57BL/6 mice as vaccinated groups. After 4 weeks, the vaccinated and non-vaccinated control groups were challenged with the same number of viable melanoma cells from paired models.

Tumor size and body weight were measured twice per week and the endpoint was established as the occurrence of one of the following events: (1) the tumor size reached 2000 mm^3^; (2) tumor became ulcerated and/or (3) the mouse showed moribund status or sickness behavior.

### Immunohistochemistry (IHC)

For tumor-infiltrating lymphocytes (TIL) analysis of M4 cell line-derived allografts, tumors were measured twice per week and collected between days 32 and 39 post-implantation when growth kinetics distinguished responders and non-responders to αCTLA-4. Non-responder tumors measured between 190-400mm^3^, and responder tumors measured between 20-125 mm^3^ at the time of sample collection. IgG control tumors were collected at day 39 post-injection (except one tumor collected at day 32). IgG control tumors measured between 400-1200 mm^3^ at the time of sample collection.

Formalin-fixed and paraffin-embedded 5 μm sections of GDAs and CLDAs from each mouse model were stained with hematoxylin and eosin (HE), DCT (Pep8h^62^), TYRP1 (clone Ta99, Invitrogen, #MA1-25303), SOX10 (clone EP268, Cell Marque, 383R-15) or CD3 (clone CD3-12, BioRad, MCA1477) antibodies using standard immunohistochemistry methods. Antigen retrieval was performed in a pressure cooker using target retrieval buffer pH6 (Dako, S1699). Protein Blocking reagent (Dako, X0909) and Bloxall blocking solution (Vector, SP-6000-100) were used to block nonspecific proteins and endogenous peroxidase and alkaline phosphatase activities. The antibodies were incubated overnight at 4°C except for SOX10, which was incubated for 30 minutes at RT, in a humidity chamber. Antibody detection was performed using Impress AP Reagent anti-rabbit Ig (Vector, MP-5401) or Impress AP Reagent anti-rat IgG (Vector, MP-5444) and ImPACT Vector Red (Vector, SK-5105). Slides were counterstained with Mayer’s Hematoxylin (Electron Microscopy Sciences, 26043-06) and GIEMSA (New Comer Supply, 1120A). Digital image files of slides were visualized using Aperio ImageScope (Leica Biosystems). Image analysis of IHC slides labeled with ImPACT Vector Red was performed using an Aperio color deconvolution algorithm.

### T cell activation assays and flow cytometry

For the T cell activation assays, M1, M2 and M4 cells were transduced with a lentiviral vector for the expression of a 9aa peptide from the mouse DCT protein^46^ fused to the C-terminus of eGFP protein but separated by a 2aa linker to ensure cleavage and presentation by MHC-I^47^. GFP^+^ cells were sorted in a BD FACSAria Fusion device and *GFP-Dct* expression was confirmed by mRNA reverse transcription (SuperScript™ III Reverse Transcriptase, ThermoFiser Scientific, 18080044) and quantitative PCR (Kappa Biosystems, KK4603) (RT-qPCR). Naïve CD8^+^ T cells from C57BL/6 splenocytes were sorted using anti-mouse CD8a particles (BD Biosciences, 551516) and cultured under CD3, CD28 and IL-2 stimulation for 2 days. CD8^+^ T cells were transduced with recombinant TCR Vβ3 for the recognition of the 9 aa DCT peptide (DCT-Vβ3) or Thy1.1 retroviral vectors and left resting (without CD3 and CD28 stimulation) for one day. Transduction efficiency was confirmed by flow cytometry using BD FACSCanto II device. Recombinant Vβ3^+^ and Thy1.1^+^ T cells were co-cultured with the melanoma cells from each model in a 1:1 ratio. After 24-hour co-culture, the concentration of Interferon-γ (IFNγ) in the media was measured by ELISA (Invitrogen, 88-7314). The T cells from 24-hour co-cultures were harvested, stained with CD8a-APC (clone 53-6.7, eBiosciences, 17-0081-82), CD137-PE (41BB, clone 17B5, BioLegend, 106106), Vβ3-FITC (clone KJ25, BD Pharmigen, 553208) and CD90.1 (Thy1.1)-PECy7 (clone HIS51, eBiosciences, 25-0900-82) antibodies and analyzed by flow cytometry using a BD LSR Fortessa SORP I device. All flow cytometry data were analyzed using FlowJo v10 software.

For the validation of MHC-I-related gene expression in the cell lines from the four models, cells were cultured for 24 hours in media containing 20ng/mL of IFNγ. MHC-I-related gene expression was analyzed by RT-qPCR. Cell were stained with H-2Kd-APC antibody (clone AF6 88.5, BioLegend, 116518) and analyzed by flow cytometry using BD FACSCanto II device.

### Phenotypic analysis of tumor immune infiltrate

For immunophenotyping, 1.0×10^6^ melanoma cells from each model were implanted subcutaneously into C57BL/6 mice. When tumors reached 100-300 mm^3^, mice were euthanized and tumors harvested and processed as previously described^63^. Briefly, tumors were weighed, mechanical disrupted and incubated in RPMI medium containing 1mg/ml of DNase I (Roche, cat 10104159001), 200U of collagenase IV (Gibco, cat 17104-019), and 0.5% FCS for 40 min at 37°C. Red blood cells were lysed using ACK buffer and cells resuspended in 2mM EDTA PBS and stained with LIVE/DEAD Fixable Dead Cell Stain Kit (Invitrogen) for 15 minutes at 4°C. Next, cells were stained with the corresponding antibody cocktail prepared in Brilliant stain buffer (BD Biosciences). Fc receptors were blocked with anti-CD16/32 antibody (2.4G2, Bioxcell) and the following anti-mouse antibodies were used: CD45.2 (clone 104), CD4 (GK1.5), CD62L (MEL-14), CD223/Lag3 (C9B7W), CD279/PD1 (RMP1-30), TCRβ (H57-597), NK1.1 (PK-136), TIGIT (1G9), Ly6G (1A8), CD64 (X54-5/7.1), CD274/PDL1 (MIH5), CD24 (M1/69), Ly6C (AL-21), CD135 (A2F10.1), SiglecF (E50-2440), purchased from BD Biosciences; CD127 (A7R34), CD8a (53-6.7), Ly6G (1A8), CD44 (IM7), CD103 (2E7), CX3CR1 (SA011F11), ICAM1 (YN1/1.7.4), NK1.1 (PK136), CD206 (C068C2), F4/80 (BM8) purchased from BioLegend; Tim3 (RMT3-23), F4/80 (BM8), CD19 (eBio1D3), TCRgd (eBioGL3), CD25 (PC61 5.3), KLRG1 (2F1), MHC II (M5/114.15.2), TCRβ (H57-597), Ter119 (TER-119), CD19 (eBio1D3), CD11c (N418), CD11b (M1/70), purchased from eBioscience. When biotinylated antibodies were used, cells were subsequently incubated with fluorochrome-conjugated streptavidin. Following surface staining, cells were fixed and permeabilized using Foxp3/Transcription Factor Staining Buffer Kit (eBioscience) according to the manufacturer’s instructions and intracellularly stained with a combination of the following anti-mouse antibodies: Foxp3 (FJK-16s) and CD3 (145-2C11) from eBioscience, CD68 (FA-11) from BioLegend. Samples were acquired in a FACS Symphony A5 running under BD FACSDiva software (BD Biosciences) and data was analyzed with FlowJo software (v10.6.0). Further analysis was done with R package Cytofkit2 (Jinmiao Chen’s Lab)^64^. tSNE algorithm was run with a concatenation of 5 samples per tumor type and the four tumor types simultaneously. For cluster analysis, Rphenograph was selected in the same package (k=30) and the clusters were lately annotated based on the expression level of the markers of interest. Clusters representing the populations of interest were selected and quantified in FlowJo. Frequencies from total leukocytes (Live CD45^+^ cells) were determined and absolute numbers per mg of tumor were calculated.

### Spectral karyotyping (SKY)

Cells from the four melanoma models were arrested in metaphase by incubation with 10μg/ml of Colcemid (15210-040, KaryoMax^®^ Colcemid Solution, Invitrogen, Carlsbad, CA, USA) 3 hours prior to harvest. Cells were collected and treated with hypotonic solution (KCL 0.075M,6858-04, Macron Chemical) for 15 minutes at 37°C and fixed with methanol:acetic acid 3:1. Slides were prepared and aged overnight for use in SKY analysis.

The metaphases were hybridized with the 21-color mouse SKY paint kit (FPRPR0030, ASI) according to the manufacturer’s protocol ^65^. Hybridization was carried out in a humidity chamber at 37°C for 16 hours. The post-hybridization rapid wash procedure was used with 0.4XSSC at 72°C for 4 minutes. Detection was carried out following the ASI manufacturer’s protocol. Spectral images of the hybridized metaphases were acquired using Hyper Spectral Imaging system (Applied Spectral Imaging Inc., CA) mounted on top of an epi-fluorescence microscope (Imager Z2, Zeiss). Images were analyzed using HiSKY 7.2 acquisition software (Genasis, Applied Spectral Imaging Inc., CA). G-banding was simulated by electronic inversion of DAPI counterstaining. An average of 10-15 mitoses of comparable staining intensity and quality was examined per cell line and compared for chromosomal abnormality. The karyotype was determined by comparison to the standard ideogram of banding patterns for mouse chromosomes^66^.

### Whole-exome sequencing

Mouse genomic DNA concentration was measured using the picogreen assay. 200ng of DNA was sheared by Covaris Instrument (E210) to ^∼^150-200 bp fragments. Shearing was done in a covaris snap cap tube (microTUBE Covaris p/n 520045) with the following parameters: duty cycle, 10%; intensity, 5; cycle burst, 200; time, 360 sec at 4°C. Samples quality assessment/size was validated by Bioanalyzer DNA-High sensitivity kit (Agilent # 5067-4626).

Agilent SureSelectXT library prep ILM Kit (Agilent Technologies, #G9611A) was used to prepare the library for each sheared mouse DNA sample. DNA fragment’s ends were repaired, followed by Adenylation of the 3’ end and then ligation of paired-end adaptor. Adaptor-ligated libraries were then amplified (Pre-capture PCR amplification: 98°C 2 minutes, 10 cycles; 98°C 30 seconds; 65°C 30 seconds; 72°C 1 minute; then 72°C 10 minutes, 4°C hold) by Herculase II fusion enzyme (Agilent Technologies, #600679). After each step DNA was purified by Ampure XP beads (Beckmann Coulter Genomics #A63882). DNA Lo Bind Tubes, 1.5-mL PCR clean (Eppendorf # 022431021) or 96 well plates were used to process the samples.

Samples were analyzed by bioanalyzer using DNA-1000 kit (Agilent #5067-1504). Concentration of each library was determined by integrating under the peak of approximately 225-275bp. Then each gDNA library (^∼^750-1000 ng) was hybridized with biotinylated mouse specific capture RNA baits (SureSelectXT Mouse All Exon, catalog #5190-4641, 16 reactions) in the presence of blocking oligonucleotides. Hybridization was done at 65°C for 16 hours using SureSelectXT kit reagents. Bait-target hybrids were captured by streptavidin-coated magnetic beads (Dynabeads MyOne Streptavidin T1, Life Technologies, #6560) for 30 minutes at room temperature. Then after a series of washes to remove the non-specifically bound DNA, the captured library was eluted in nuclease free water and half volume was amplified with individual index (Post-capture PCR amplification: 98°C 2 minutes; 10 cycles, 98°C 30 seconds; 57°C 30 seconds; 72°C 1 minute; then 72°C 10 minutes, 4°C hold). The Bioanalyzer High sensitivity kit was used to validate the size of the libraries prior to sequencing.

### Single nucleotide variant (SNV) analysis

All analyses were carried out on NIH biowulf2 high performance computing environment. Statistical analysis was carried out in R environment.

Fastq sequence reads were mapped to the mouse reference genome mm10 with BWA or Bowtie. Single nucleotide variants (SNV) were identified using samtools mpileup or GATK HaplotypeCaller. Mouse germline single nucleotide polymorphisms (SNPs) were filtered out the Sanger database for variants identified from whole genome sequencing of 36 mouse strains (ftp://ftp-mouse.sanger.ac.uk/current_snps/mgp.v5.merged.snps_all.dbSNP142.vcf.gz). Variants with a Phred-scaled quality score of <30 were removed. Variants that are present in normal spleen samples (in-house collection) were also removed. Variants were annotated with Annovar software to identify non-synonymous mutations.

For mutation signature analysis, the 5 prime and 3 prime nucleotide sequences flanking the mutations were retrieved from the mm10 reference genome using bedtools getfasta. The frequency of 96 trinucleotides (6 substitutions multiplied by 16 combinations of 5 and 3 prime nucleotides) was computed for each sample with an in-house R script. Cosmic Signatures^41^ were identified using R package deconstructSigs.

For clustering analysis with TCGA samples, we used 20 genes that are most frequently mutated in human melanoma studies ^43^. The samples (the cell lines or GDAs from the 4 models or TCGA melanomas) that have a mutation from the 20-gene list were coded as 1 for that gene whereas the samples without mutation were coded as 0 for that gene. The heatmap was generated using R package ComplexHeatmap.

### RNA sequencing

Between 100ng to 1ug of total RNA was used to capture mRNA with oligo-dT coated magnetic beads. The mRNA was fragmented, and then a random-primed cDNA synthesis was performed. Alternatively, total RNA from GDA2s and CL4 was processed to remove ribosomal RNA (rRNA) using biotinylated, target-specific oligos combined with Ribo-Zero rRNA removal beads. The RNA was fragmented into small pieces and the cleaved RNA fragments were copied into first strand cDNA using reverse transcriptase and random primers, followed by second strand cDNA synthesis using DNA Polymerase I and RNase H. The resulting double-strand cDNA was used as the input to a standard Illumina library prep with end-repair, adapter ligation and PCR amplification being performed to obtain a sequencing ready library. The final purified product was then quantitated by qPCR before cluster generation and sequencing.

### Differential gene expression analysis

The sequence reads in fastq format were aligned to the mouse reference genome mm10 using STAR^67^ and mapped sequence reads were normalized by RSEM^68^ to obtain the fragments per kilobase of transcript per million (FPKM) per gene. Pairwise differential expression of the cell line-derived allografts from the four melanoma models was performed using DESeq2^69^. For untreated melanomas, the 20% most variable genes among the differentially expressed (DE) genes between the models (M1 vs. M3, M1 vs. M4, M2 vs. M3 and M2 vs. M4) were categorized using Ingenuity pathway analysis (IPA, QIAGEN). For αCTLA-4 vs. isotype control comparisons, genes with FDR<0.1 and Fold Change>1.5 were considered significant.

### CIBERSORT and Gene set enrichment analysis

The normalized counts of the RNA sequencing data from the cell line-derived allografts were analyzed by DESeq2 to compare between models and then the results were further analyzed by Gene set enrichment analysis (GSEA) ^70^. For each comparison between two models, we identify enriched “Hallmark” pathways and considered those significant ones with a FDR<0.05.

To evaluate the immune cell population abundances in our models, we analyzed the gene expression data of the untreated CLDAs from each model using the CIBERSORT tool ^53^.

### TIDE score prediction

The gene expression data of the untreated CLDAs from each model were analyzed by Tumor Immune Dysfunction and Exclusion (TIDE) tool using TIDE web application at http://tide.dfci.harvard.edu/. We used a cutoff value of 0 to predict the response of each tumor.

### Melanocytic Plasticity signature (MPS) development

Based on the expression of untreated cell line-derived tumors from the 4 mouse models, the differentially expressed (DE) genes comparing M1 and M2 vs. M3 and M4 were identified by t-test on the RPKM data and by DESeq2 on the RNASeq count data. The DE genes were ranked using three criteria: 1) by p-values from the t-test, 2) by log2FoldChange from DESeq2 and 3) by FcPv=-log2FoldChange*log10(padj) from DESeq2. For each of the three gene lists ranked by the three approaches, we started from the two most upregulated genes and stepwise adding next most upregulated gene to obtain a list of gene sets and using the Van Allen data set ^4^ to find the best signature to predict patient response based on Fisher’s exact test. Similarly, we found the best signatures from downregulated genes. We removed the signatures that were not significant using the Fisher’s exact test. Combining the significant signatures, we obtained a 45-gene signature comprised of 33 genes up and 12 genes downregulated in M1/M2. To calculate MPS scores the expression of the 45 genes in RPKM was weighted by 1 for upregulated genes and −1 for the downregulated genes and added.

The 45 genes in the signature were categorized by IPA analysis.

### Statistical analysis

For *in vivo* therapeutic studies, tumors were measured independently by an animal technician, and the size (V) was calculated as following: V = 0.5*L*W^2^, in which L is the longer diameter and W is the shorter diameter. The survival time of the mice was defined as the duration from tumor implantation to the occurrence of one of the following events: (1) the tumor reached 2000 mm^3^; (2) tumor became ulcerated; or (3) the mouse showed moribund status or sickness behavior. The Kaplan-Meier analysis was performed for survival of treatment groups using GraphPad Prism, and significance is analyzed by the Log-rank (Mantel-Cox) test. For vaccination studies, tumor onset time was defined as the post-implantation time when the tumor was observed to be palpable independently by an animal technician.

The Mann-Whitney test was used for the comparison between two groups using GraphPad Prism. For phenotypic analysis of tumor immune infiltrate in the four models, Kruskal-Wallis test followed by Dunn’s multiple comparisons test was performed in Prism 8 (v8.1.2, GraphPad Software). For survival analysis of Van Allen^4^ and Hugo-Riaz^8,19^ data sets, the Cox proportional hazard (Cox-PH) models were estimated from the patient survival data and used to compute hazard ratios.

## CONTACT FOR REAGENTS AND RESOURCE SHARING

Further information and requests for resources and reagents should be directed to and will be fulfilled by the Lead Contact, Glenn Merlino. Requests for the reagents derived from mouse models (alleles, tumor tissues, cell lines, etc.) and DNA constructs developed in this study will require Material Transfer Agreement per NIH guideline.

WES raw data of the GDAs and cell lines from the four models and RNA sequencing raw data of the CLDAs from the four models will be deposit on GEO.

## ACKNOWLEDGMENTS

We thank: Dr. Timothy A. Chan (Memorial Sloan-Kettering Cancer Center, New York, NY, USA) for suggestions and comments; Dr. Thomas Tüting (University of Magdeburg, Magdeburg, Germany), Dr. Martin McMahon (University of Utah, Salt Lake City, UT, USA) and Dr. Marcus Bosenberg (Yale School of Medicine, New Haven, CT, USA) for mouse reagents; Dr. Eliezer Van Allen (Dana-Farber Cancer Institute, Boston, MA, USA) for sharing clinical data; Sandra Burkett (Molecular Cytogenetics Core Facility, MCGP, National Cancer Institute-Frederick, Frederick, MD, USA) for SKY analysis; Dr. Christophe Redon (National Cancer Institute, NIH, Bethesda, MD, USA) for assistance with gamma irradiation; Dr. Yanis Boumber (Fox Chase Cancer Center, Philadelphia, PA, USA) and Dr. Christine Alewine (National Cancer Institute, NIH, Bethesda, MD, USA) for manuscript revision and editing. This research was supported in part by funds from the NIH intramural research program and a FLEX Synergy Award from the NCI Center for Cancer Research. In addition, an NCI Director’s Innovation Award to EPG helped support this project.

## AUTHOR CONTRIBUTIONS

Conceptualization, E.P-G., C-P.D. and G.M.; Methodology, R.EM., H.M., S.K.V. and C-P.D.; Software, H.H.Y. and M.P.L.; Validation, E.P-G., H.H.Y., M.P.L. and C-P.D.; Formal Analysis, E.P-G., H.H.Y., R.EM., H.M., S.K.V., A.M., M.P.L. and C-P.D.; Investigation, E.P-G., R.E.A., R.EM., S.K.V., C.G-C., S.C., A.F., A.J.I., A.K., W.H., R.S.L. and C-P.D.; Data Curation, H.H.Y., A.M. and M.P.L.; Writing – Original Draft, E.P-G., C-P.D. and G.M.; Writing – Review and Editing, E.P-G., H.H.Y., R.EM., H.M., S.K.V., N.P.R., T.VD., S.K.S., R.S.G, Z. W-O., C-P.D. and G.M.; Visualization, E.P-G., H.H.Y., R.E.A., M.P.L. and C-P.D.; Supervision, S.K.S., R.S.G., Z.W-O., M.P.L., C-P.D. and G.M.; Project Administration, E.P-G., R.EM. and C-P.D.; Funding Acquisition, G.M.

