## Supplementary figures and tables for "Multi-model preclinical platform predicts clinical response of melanoma to immunotherapy"

| Melanoma Model | M1 | M2 | M3 | M4 |
|---|---|---|---|---|
| Mouse strain | Albino C57BL/6 | C57BL/6 | C57BL/6 | C57BL/6 |
| Genotype | $Braf^{CA/+}$; $Pten^{flox/+}$; $Cdkn2a^{flox/+}$; $Tyr\text{-}Cre^{ERT2\text{-}tg}$ | $Braf^{CA/+}$; $Cdkn2a^{flox/+}$; $Tyr\text{-}Cre^{ERT2\text{-}tg}$; $Hgf^{tg}$ | $Cdk4^{R24C}$; $Hgf^{tg}$ | $Hgf^{tg}$ |
| Carcinogen | UV | UV | DMBA | UV |

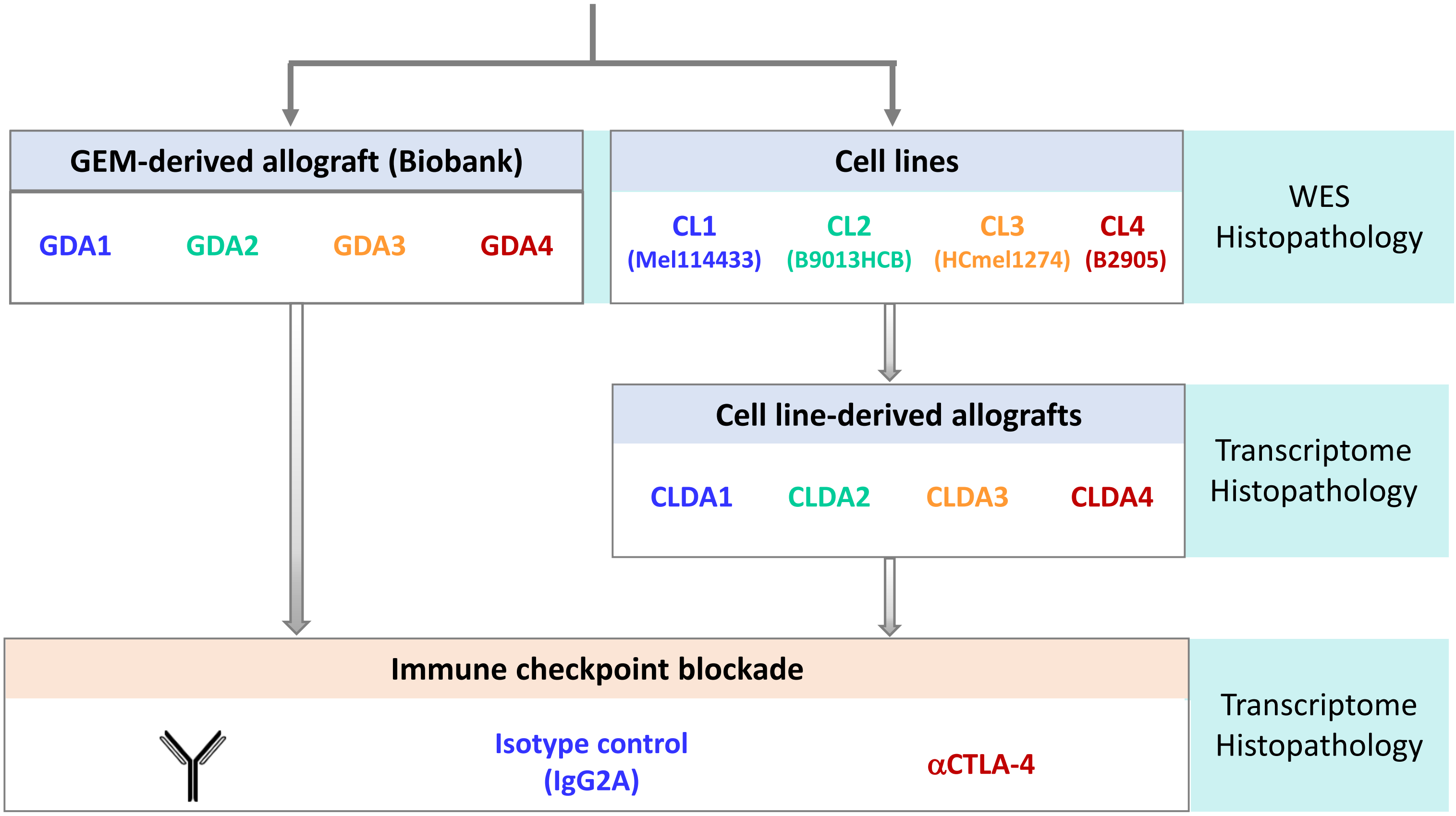


M: Melanoma model
GDA: Genetically engineered mouse (GEM)-derived allograft
CL: Cell line
CLDA: Cell line-derived allograft
WES: Whole exome sequencing

**Supplementary Fig. 1: Study design and sample processing.** Melanomas were induced in four genetically engineered mice (GEM) harboring the indicated genetic modifications by ultraviolet (UV) radiation or 7,12-Dimethylbenz(a)anthracene (DMBA) topical administration at postnatal day 3 and activation of CreERT2 at day 7. Fragments from each melanoma were expanded in C57BL/6 syngeneic mice and viably archived at low passage to generate a GEM-derived allograft (GDA) biobank. A cell line (CL) from each model was isolated. GDA and cell line-derived allografts (CLDA) were implanted in C57BL/6 mice and treated with CTLA-4 blocking antibody (αCTLA-4) or isotype control. GDAs, CLDAs and cultured cell lines from each model were processed in triplicates for whole exome sequencing (WES), RNA sequencing and/or histopathology.

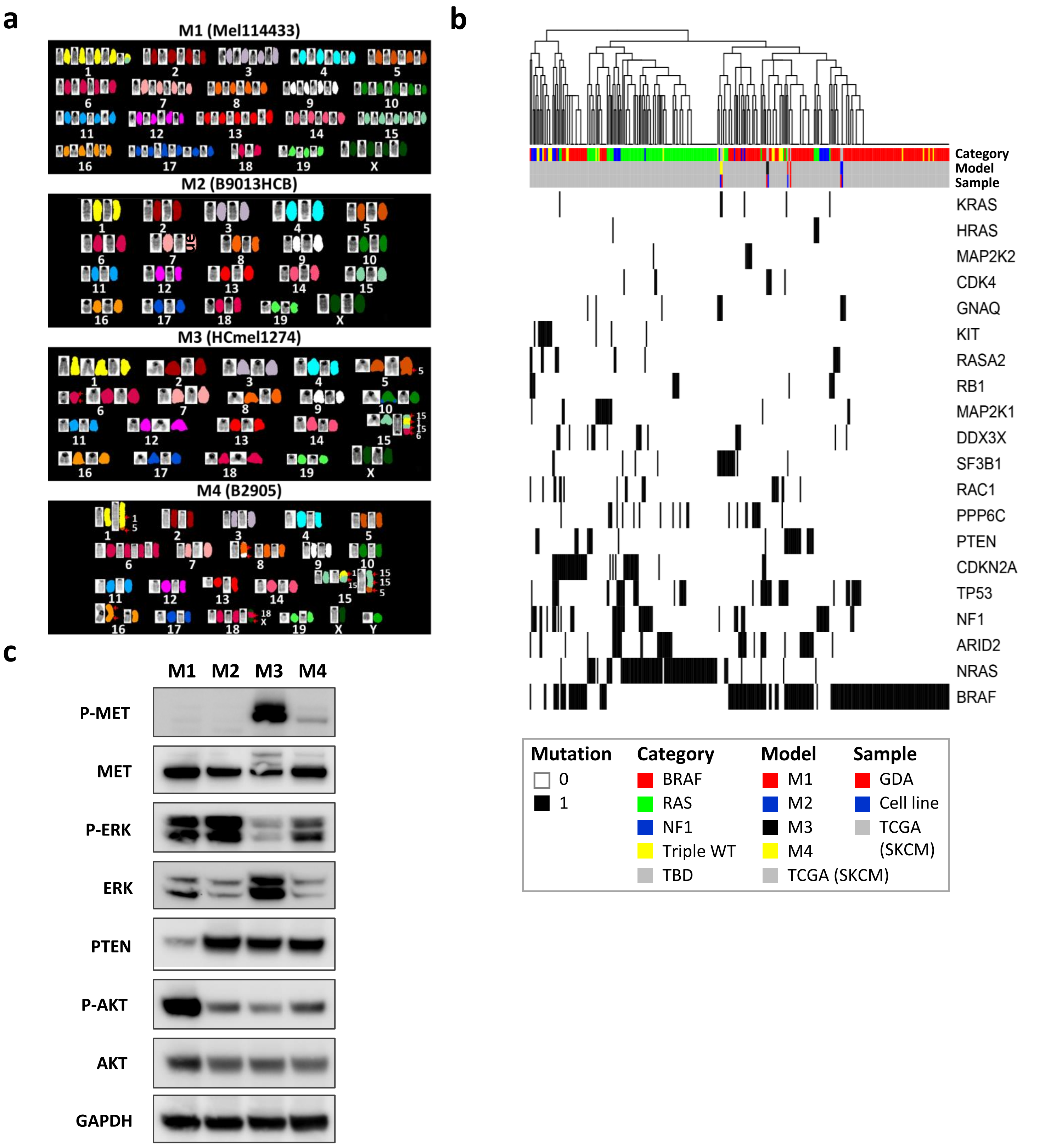


**Supplementary Fig. 2: Generation of four mouse models representative of diverse human melanomas. a,** Representative images showing chromosomal duplications and translocations analyzed by spectral karyotyping (SKY) of the four melanoma cell lines. See also **Supplementary Table 2**. **b,** GDAs and cell lines derived from the four mouse models were clustered accordingly to their mutation profiles with TCGA patient samples from different mutation categories (i.e. BRAF, NRAS or NF1 mutants, or triple-wildtype [36,43]). **c,** Immunoblot showing MAPK and PI3K activation by the expression of the indicated proteins in the four melanoma cell lines.

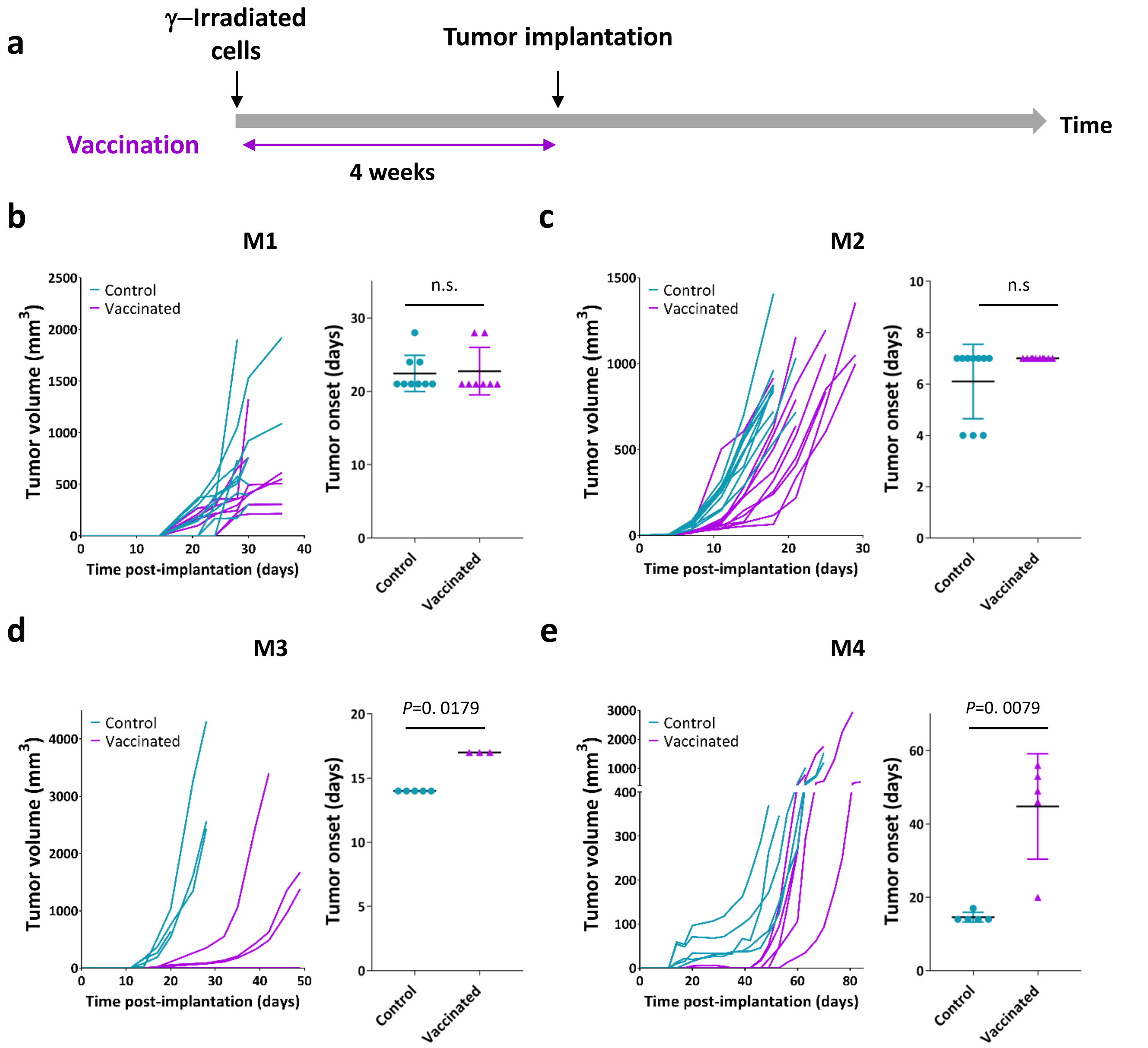


**Supplementary Fig. 3: Tumor immunogenicity correlates with αCTLA-4 response in the melanoma models. a,** Schematic of the *in vivo* study design. 1.0x$10^6$ γ-irradiated melanoma cells from each model were s.c. injected into C57BL/6 mice as vaccinated groups. After 4 weeks, the vaccinated and non-vaccinated control groups were challenged with the same number of viable melanoma cells from paired models. **b-e,** Tumor growth curves (left panels) and tumor onset (right panels) of vaccinated (magenta) and control (light blue) groups from M1 (b, N=9 in control and N=8 in vaccinated group), M2 (c, N=10), M3 (d, N=5) and M4 (e, N=5). The time of tumor onset was considered as the first day after implantation when tumors were measurable. *P*-values from Mann-Whitney test are as indicated (b-e, right panels). n.s.: non-significant.

Supplementary Fig. 4

Pérez-Guijarro et al.

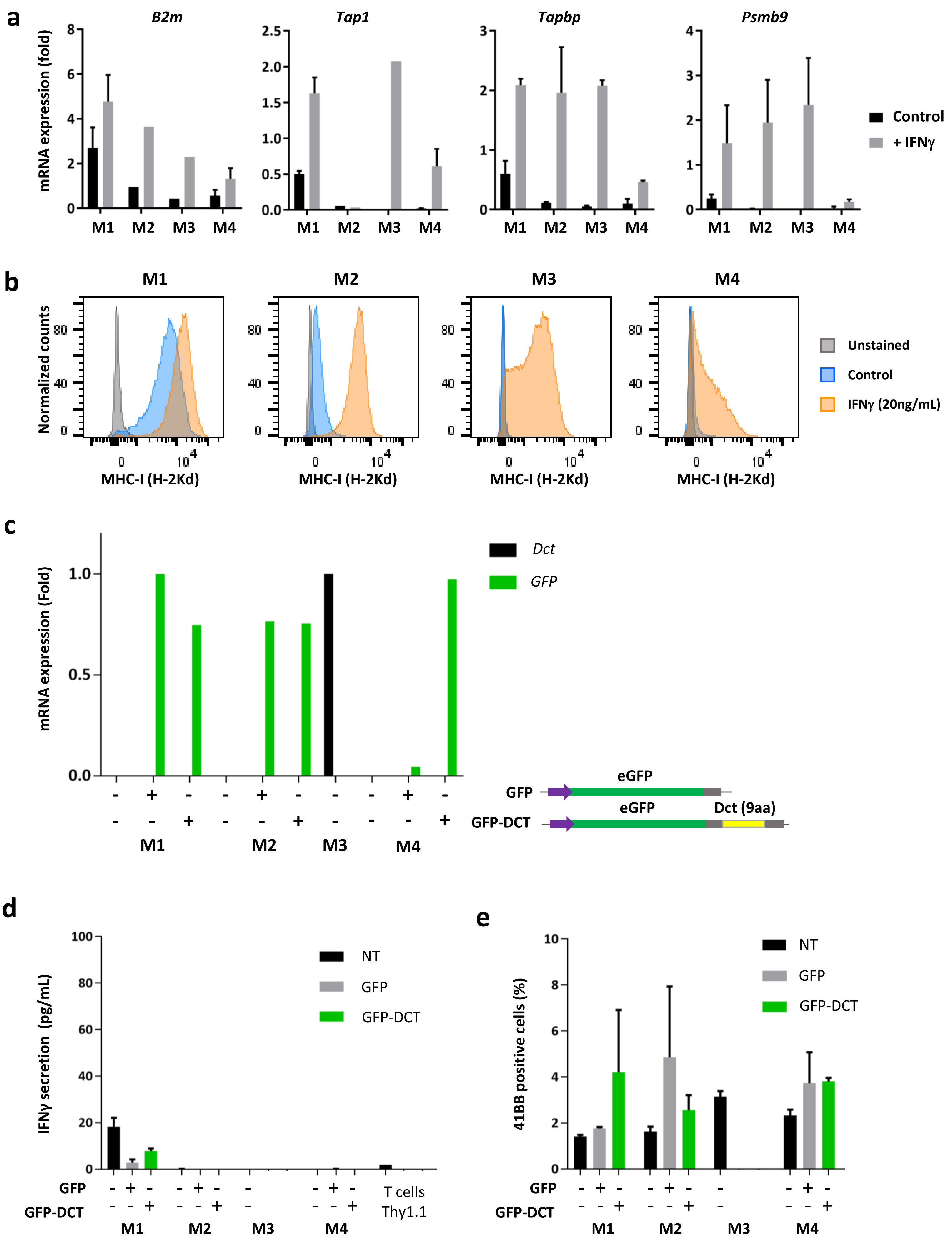

**Supplementary Fig. 4: Antigen presentation pathway is induced in the four models. a,** Validation of expression of the indicated MHC-I-related genes by RT-qPCR in cell lines from the four models. Cells were cultured for 16 hours in 20ng/mL Interferon-γ (IFNγ, grey bars) or standard culture conditions (black bars). Graph depicts the mean of 3 independent experiments and error bars represent S.E.M. **b,** Analysis of MHC-I (H-2Kd) expression by flow cytometry of IFNγ-stimulated (red line) or control (back line) cells. Unstained cells were used as negative control (grey line). **c,** Endogenous *Dct* mRNA (grey bars) and *GFP-Dct* or *GFP* (green bars) expression levels in the indicated melanoma cells measured by RT-qPCR. The 9aa DCT peptide[46] was expressed in frame with eGFP for its proper cleavage and presentation by MHC-I[47]. **d,** IFNγ secretion by Thy1.1-transduced $CD8^+$ T cells after 24-hour co-culture with the indicated melanoma cell lines measured by ELISA. **e,** Percentage of 41BB positive $CD8^+$ T cells that were transduced with Thy1.1 control vector and co-cultured with the indicated melanoma cells. Co-cultures were done in triplicates and error bars represent S.E.M. (d,e). A representative of 2 independent experiments is shown (b-e).

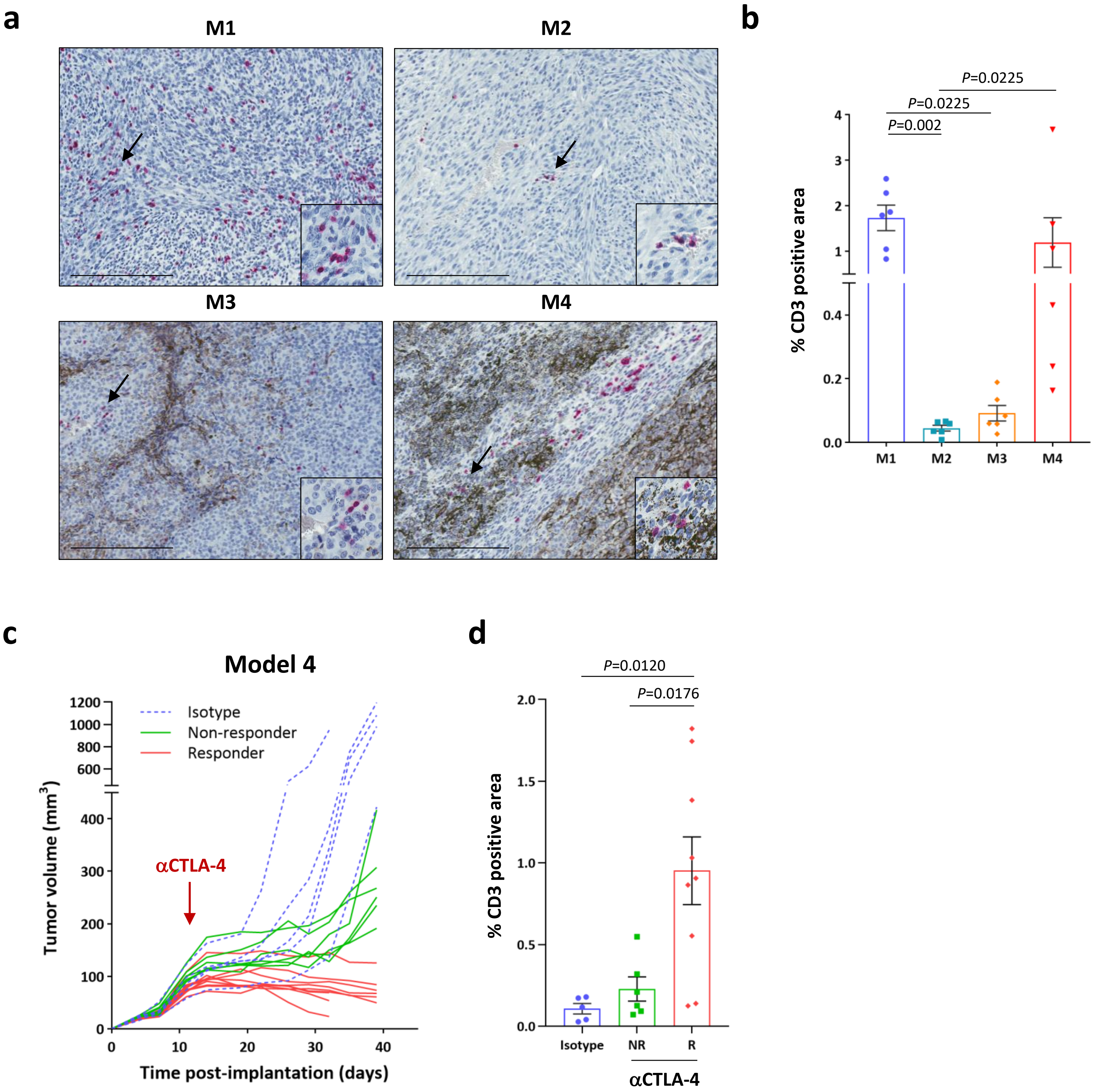


**Supplementary Fig. 5: Increased tumor-infiltrating lymphocytes are associated with αCTLA-4 response in the melanoma models. a,** Representative images of CD3 immunostaining (red) of the four melanoma models (untreated). Arrows point to the area magnified and bars represent 200 μm. **b,** Automated quantification of CD3 positive area obtained by immunohistochemistry. Whole-tumor sections from melanomas representing each model (N=6) were quantified using Aperio software. **c,** Tumor growth curves of M4 melanomas treated with αCTLA-4 (green and red lines) or isotype antibody (blue dashed line). Tumors were considered non-responders (NR, green) when their size was >150mm$^3$ and increasing for more than 2 consecutive measurements and responders (R, red) when their size was <120mm$^3$ and decreasing for more than 2 consecutive measurements. **d,** Percentage of CD3 positive area in whole-tumor sections from M4 treated with αCTLA (NR, green bar and R, red bar) or isotype antibody (blue bar). Data is depicted as the mean and error bars represent S.E.M (N=5 for isotype control, N=6 for NR and N=9 for R). Mann-Whitney test *P*-values are indicated.

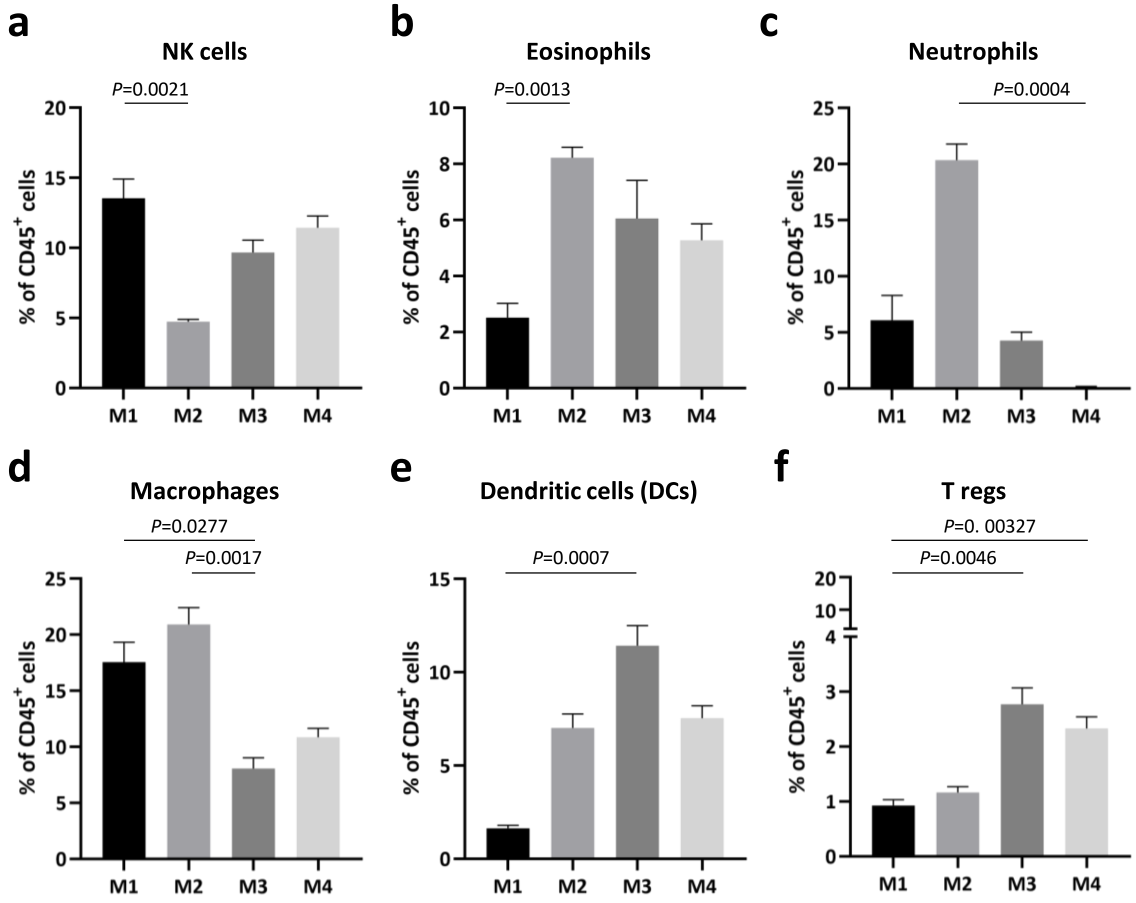


**Supplementary Fig. 6: High parametric flow cytometry analysis of the intratumoral immune cell populations from the four melanoma models. a-f,** Percentage of intratumoral NK cells identified as $CD3^-$ $NK1.1^+$ (a), eosinophils ($CD11b^+$ $Ly6G^-$ $Ly6C^{int}$ $SiglecF^+$ $F4/80^+$ $CD64^-$ $CD24^+$) (b), neutrophils ($CD11b^+$ $Ly6G^+$ $Ly6C^{int}$ $SiglecF^-$ $CD24^+$) (c), macrophages ($CD11b^+$ $CD68^{hi}$ $CD64^+$ $F480^+$ $Ly6C^-$ $Ly6G^-$ $SiglecF^-$ $CD24^-$) (d), dendritic cells ($Ly6G^-$ $SiglecF^-$ $CD11c^+$ MHC-$II^+$ $CD64^-$ $CD24^{int/hi}$ $CD135^+$) (e) and Tregs ($CD3^+$ $TCR\beta^+$ $CD4^+$ $CD25^+$ $FoxP3^+$) (f) in the four models. Data from a representative of two experiments is depicted as the mean (N=5) and error bars represent S.E.M. Kruskal-Wallis test *P*-values adjusted for multiple comparisons are indicated (a-f). See also **Supplementary Table 4** for the gating description.

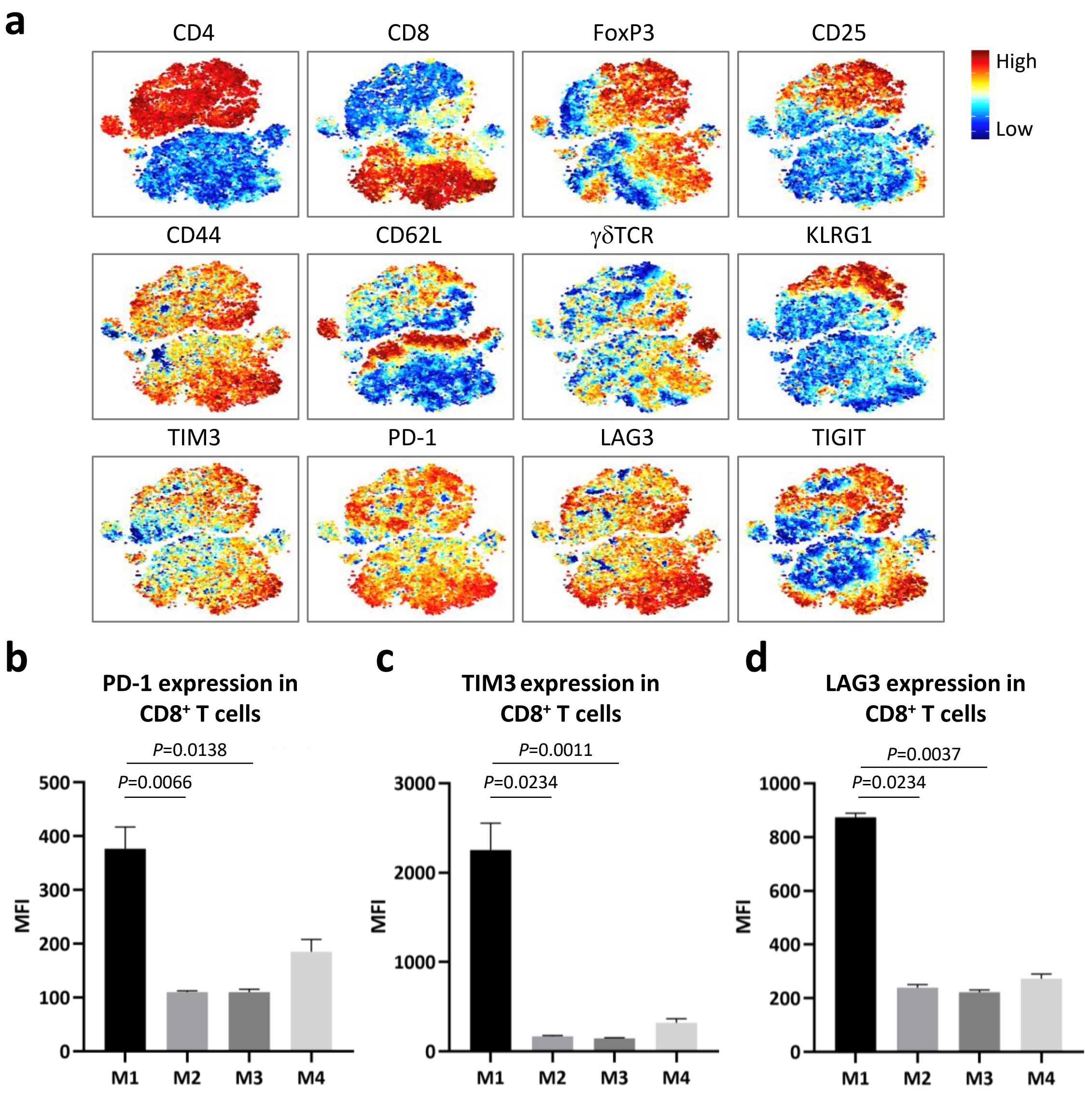


**Supplementary Fig. 7: Expression of T cell exhaustion markers in the four melanoma models. a,** Expression of the indicated markers per intratumoral $CD3^+$ T cell from the four models analyzed by t-stochastic neighbor embedding (t-SNE). **b-d,** Expression of PD-1 (b), TIM3 (c) and LAG3 (d) in $CD8^+$ T cells as the mean fluorescence intensity (MFI) from flow cytometry. Data from a representative of two experiments is depicted as the mean (N=5) and error bars represent S.E.M. Kruskal-Wallis test *P*-values adjusted for multiple comparisons are indicated (b-d). See also **Supplementary Table 4** for the gating description.

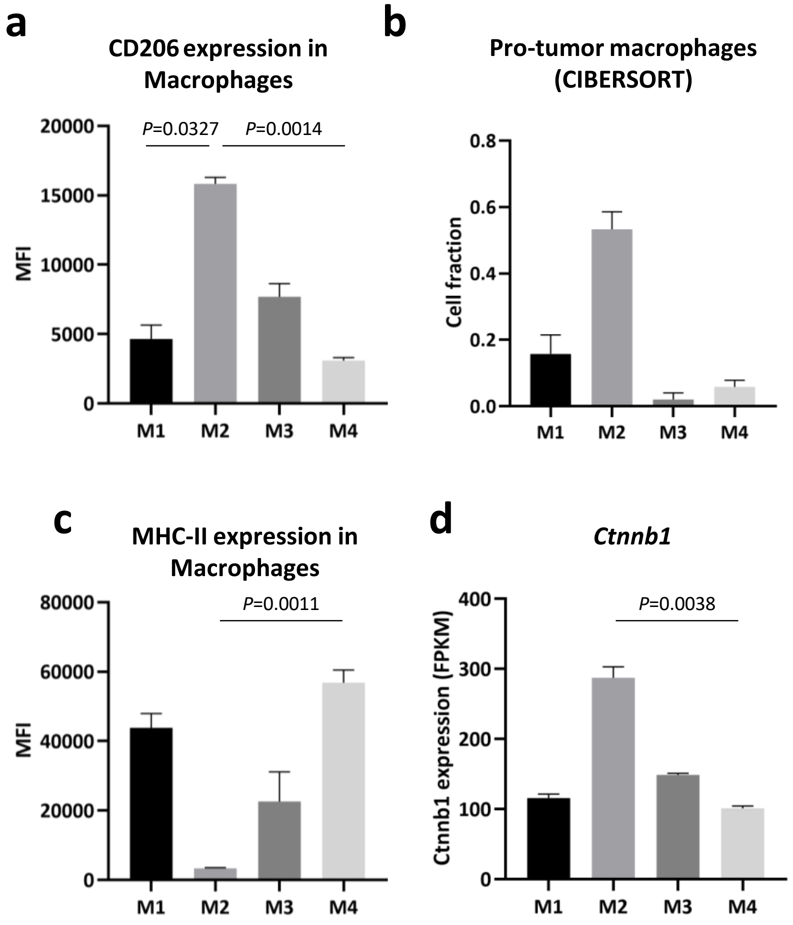


**Supplementary Fig. 8: Intratumoral immune cell populations contributing to the T cell dysfunction and exclusion profiles in the resistant models. a,c** Expression of CD206 (a) and MHC-II (c) in the intratumoral macrophages of the four models measured by flow cytometry (MFI: mean fluorescence intensity). See also **Supplementary Table 4** for the gating description. **b,** Fraction of pro-tumor macrophages obtained by CIBERSORT[50,51] analysis of the transcriptomes of the four models (untreated, N=4). **d,** *Ctnnb1* expression from RNA sequencing analysis of the four melanoma models (untreated, N=4). Data from a representative of two experiments is depicted as the mean (N=5) and error bars represent S.E.M. (a,c). Kruskal-Wallis test *P*-values adjusted for multiple comparisons are indicated (a-d).

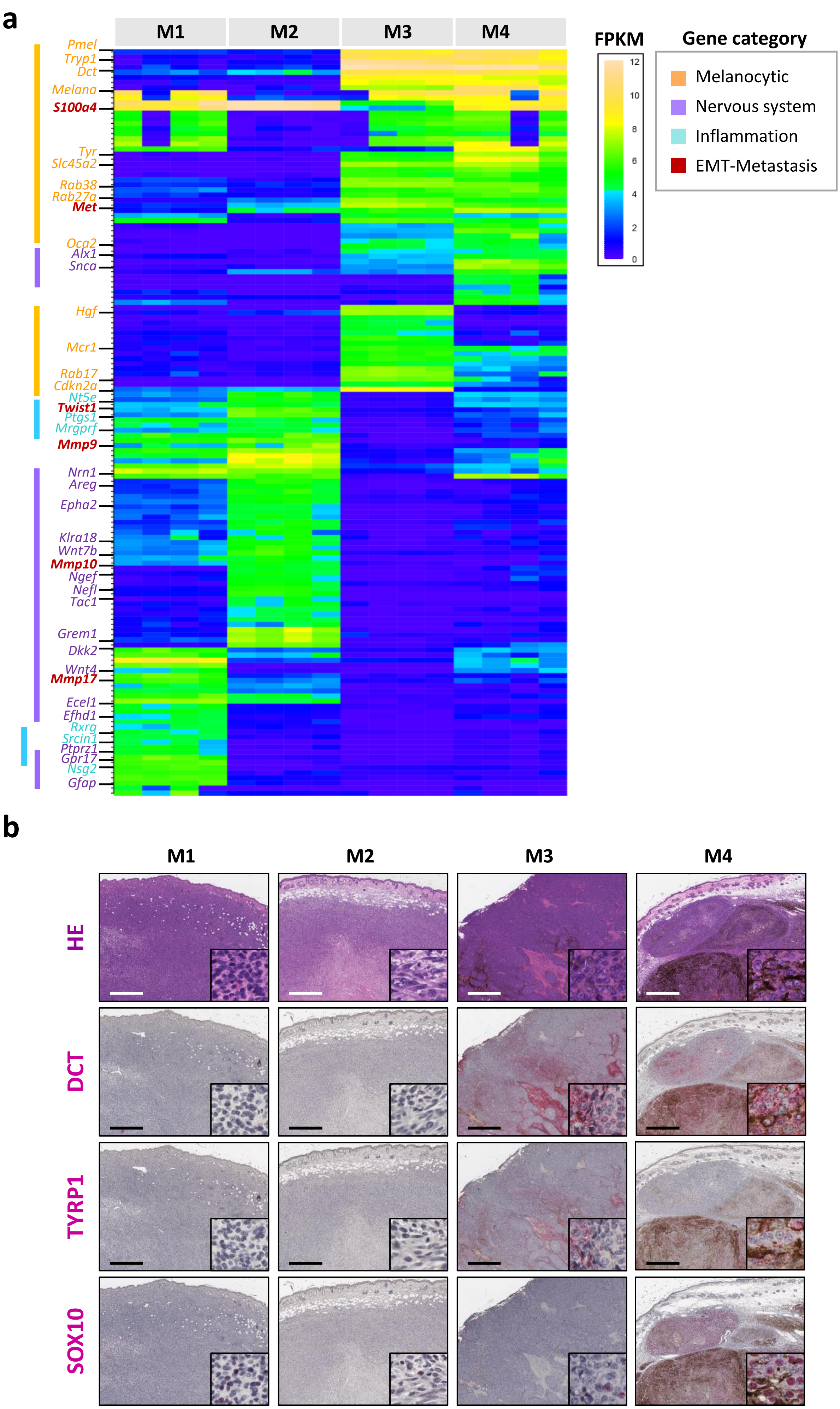
a
M1
M2
M3
M4
FPKM
12
10
8
6
4
2
0
Gene category
Melanocytic
Nervous system
Inflammation
EMT-Metastasis
Pmel
Tryp1
Dct
Melana
S100a4
Tyr
Slc45a2
Rab38
Rab27a
Met
Oca2
Alx1
Snca
Hgf
Mcr1
Rab17
Cdkn2a
Nt5e
Twist1
Ptgs1
Mrgprf
Mmp9
Nrn1
Areg
Epha2
Klra18
Wnt7b
Mmp10
Ngef
Nefl
Tac1
Grem1
Dkk2
Wnt4
Mmp17
Ecel1
Efhd1
Rxrg
Srcin1
Ptprz1
Gpr17
Nsg2
Gfap
b
M1
M2
M3
M4
HE
DCT
TYRP1
SOX10

**Supplementary Fig. 9: Distinct differentiation status of the four melanoma models. a,** Heatmap representing the 20% most variable genes among the differentially expressed genes between the models, obtained by pairwise DESeq analysis (M1 vs. M3, M1 vs. M4, M2 vs. M3, and M2 vs. M4). The heatmap depicts the fragments per kilobase million (FPKM) of the selected genes obtained from RNA sequencing of the cell line-derived allografts (N=4 per model). The highlighted genes are color-coded by their functional category annotated in Ingenuity Pathway Analysis software (IPA, "Disease and functions" categories). **b,** Representative images of hematoxylin and eosin (HE), SOX10, DCT and TYRP1 immunohistochemistry (red) in the four models. Bars represent 600 μm.

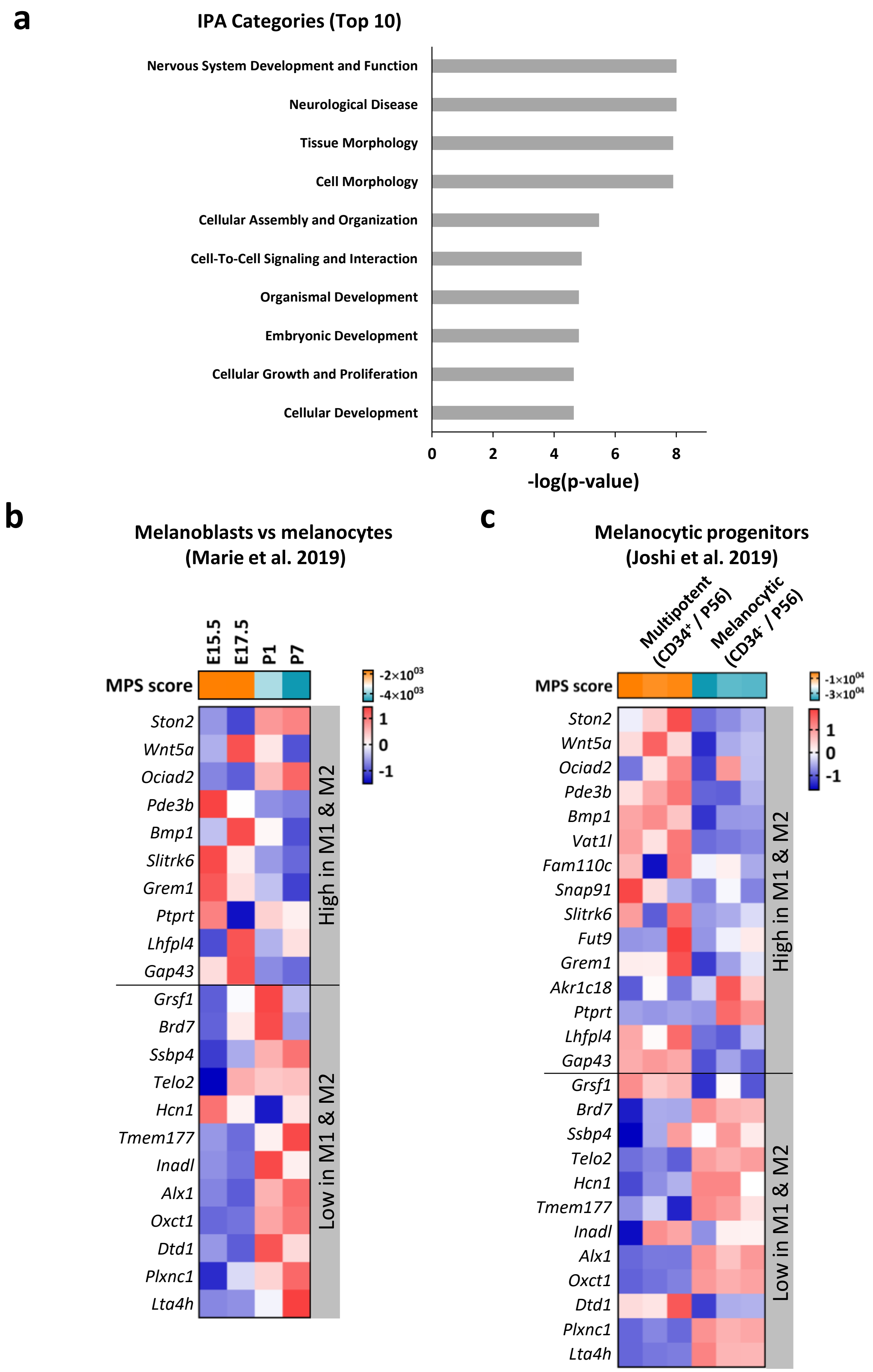


**Supplementary Fig. 10: Association of the predictive signature with melanocytic lineage differentiation. a,** Top 10 IPA Disease and Bio function categories enriched in the 45 genes of the signature. **b-c,** Expression of the melanocytic plasticity signature (MPS) genes and MPS scores of mouse melanoblasts (days E15.5 and E17.5) vs melanocytes (P1 and P7) (Marie et al., 2019) (b) and multipotent ($CD34^{+}$) vs melanocytic committed ($CD34^{-}$) melanocyte stem cells from the hair follicles of P56 mice[52] (c). Data is represented as the z-scores from RNA sequencing (b,c).

### Supplementary Fig. 11



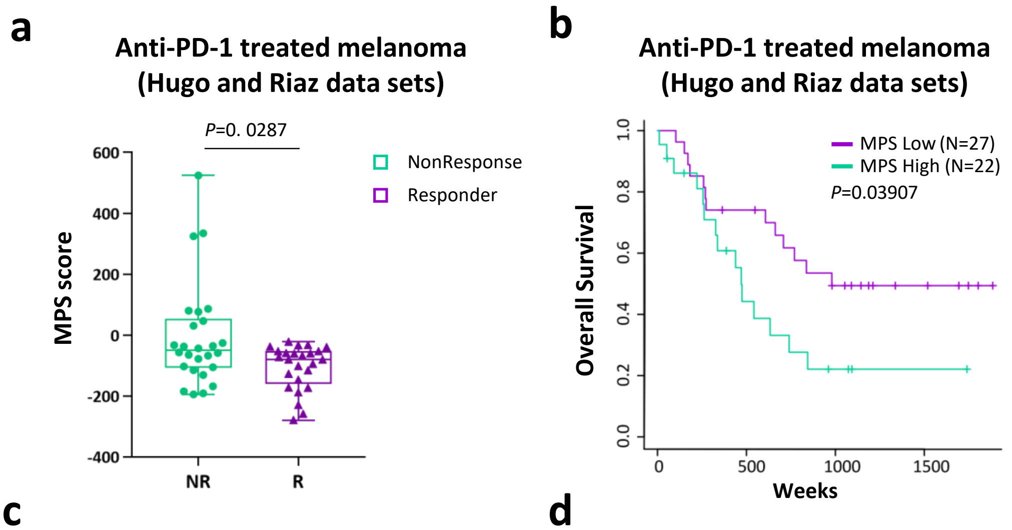


**c**

| | Van Allen | Hugo-Riaz |
|---|---|---|
| **TMB** | 0.70 | 0.63 |
| **PD-L1** | 0.62 | 0.48 |
| **TIDE** | 0.76 | 0.64 |
| **MPS** | 0.77 | 0.68 |

**AUC**

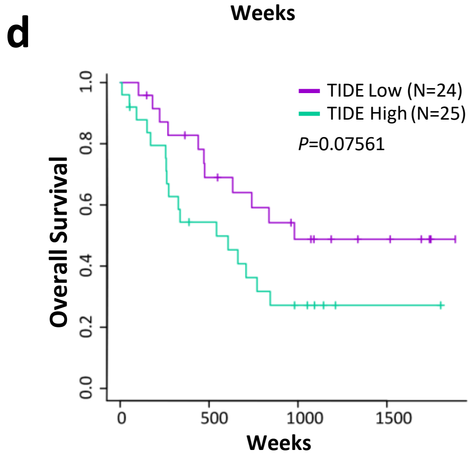


**Supplementary Fig. 11: The Melanocytic Plasticity signature (MPS) predicts patient outcome in response to αPD-1. a,** Melanocytic plasticity signature (MPS) in ipilimumab-naïve melanoma patients treated with αPD-1 (Hugo and Riaz data sets[8,19]). The box plot shows the MPS scores of baseline samples from non-responder (NR, green, N=26) and responder (R, violet, N=25) patients. Mann-Whitney test *P*-value is indicated. **b,** Kaplan-Meier curves of the overall survival of the patients in (a) accordingly to their MPS scores. **c,** Area under the ROC curve (AUC) values comparing the prediction performance of tumor mutation burden (TMB), *PD-L1* expression, TIDE and MPS scores in Van Allen[4] (left) and Hugo-Riaz[8,19] (right) data sets. **d,** Kaplan-Meier curves of the overall survival of the patients in (a) accordingly to their TIDE scores. *P*-values from the Log-rank (Mantel-Cox) test are indicated (b,d).

#### Supplementary Table 2. Spectral karyotyping (SKY) analysis of the four models cell lines

| Cell number | Number of Chromosomes | Gender | Alterations |
|---|---|---|---|
| **CL1 (MEL114433)** | | | |
| 1 | 78 | XXXX | +Del(1),-13,-13,+17 |
| 2 | 41 | XX | Del(1) |
| 3 | 83 | XXXX | +t(1;15),-4,+8,+9,+10,-13,+17,+18,-19 |
| 4 | 77 | XXXX | -8,-12,-16 |
| 5 | 82 | XXXX | +Del(1),+6,+Del(8),+10,- 12,-17 |
| 6 | 78 | XXXX | -4,-5,-8,-9,+10,+15,-16,+19 |
| 7 | 80 | XXXX | |
| 8 | 41 | XX | +Del(1) |
| 9 | 42 | XX | +T(1;15),+Del(1),+9 |
| 10 | 85 | XXX | -5,+6,+8,+10x3,+14,+15,+15,-18 |
| 11 | 83 | XXXX | +?t (1;15),+10,+13,+15,+17, |
| 12 | 84 | XXXX | +6,+8,+10,-14,+15,+15 |
| **CL2 (B9013HCB)** | | | |
| 1 | 40 | XX | |
| 2 | 40 | XX | |
| 3 | 40 | XX | |
| 4 | 40 | XX | |
| 5 | 40 | XX | |
| 6 | 40 | XX | |
| 7 | 40 | XX | |
| 8 | 40 | XX | |
| 9 | 40 | XX | |
| 10 | 40 | XX | |
| 11 | 40 | XX | Del(11),T(16;11)Looks reciprocal no loss of material |
| 12 | 40 | XX | |
| 13 | 40 | XX | |
| 14 | 40 | XX | |
| 15 | 40 | XX | |
| **CL3 (HCmel1274)** | | | |
| 1 | 42 | X | +1,Der(5),+Dic(6),+T(10;17), |
| 2 | 42 | XX | +1, Der(5),+Dic(6),T(10;17),Der(15)T(15;1;15;6) |
| 3 | 42 | XXX | +1, Der(5),+Dic(6),+T(10;17),Der(15)T(15;1;6),-19 |
| 4 | 42 | XX | +1, Der(5),+Dic(6),+T(10;17),Der(15)T(15;1;15;6)Dic(15;1;15;6) |
| 5 | 42 | XX | +1, Der(5),+Dic(6),+T(10;17),Der(15)T(15;1;6) |
| 6 | 42 | XX | +1,Der(5)add(5q),+Dic(6), T(10;17), |
| 7 | 42 | XX | +1,Del(3),Der(5),+Dic(6),+T(10;17), |
| 8 | 42 | XX | +1, Der(5),+Dic(6),+T(10;17),Dic(15)T(15;1;15;6) |
| 9 | 43 | XX | +1, Der(5),+Dic(6),+T(10;17), t(11;11), |
| 10 | 42 | XX | +1, Der(5),+Dic(6),+T(10;17),Dic(15)T(15;1;15;6) |
| 11 | 42 | XX | +1, Der(5),+Dic(6),+T(10;17),Der(15)Dic T(15;1;15;6) |
| 12 | 43 | XX | +2,Der(5),+Dic(6)(A1-A3;A3-A1), T(10;17),+17 |
| 13 | 42 | XX | +1,Del(3),Der(5),+Dic(6),+T(10;17), Der(15)T(15;1;6) |
| 14 | 41 | XX | +1,Del(3),Der(5),+Dic(6),+T(10;17), Der(15)T(15;1;6),-16 |
| 15 | 42 | XX | +1,Der(5),+Dic(6),T(10;17) |
| **CL4 (B2905)** | | | |
| 1 | 41 | XY | Der(1)?T(1;15;1),+3,-4,+6,+6,+T(8;9),-9.T(2;13), -14,T(16;16),-17 |
| 2 | 45 | XY | Der(1)T(1;5),+6,+6,+T(8;9),Del(13),+Der(15)T(15;15;5), T(1;15),T(16;16),T(18;X) |
| 3 | 44 | XY | -1,+6,+T(8;9),Del(9)x2,+T(9;9), Del(13),+Der(15)T(15;15), |
| 4 | 47 | XY | +Del(1),+3,+6,+6,+T(8;9),+Del(12),+T(15;1) |
| 5 | 45 | XY | +Del(X),Der(1)? T(1;15;1),+3,+6,+T(8;9),- 9,-13,T(2;13),T(16;16),+Der(18) |
| 6 | 45 | XY | +Del(X), Del(1),+3,+6,+8,T(9;9),-10,Del(13),T(16;16),+18 |
| 7 | 42 | X | +Der(1)? T(1;15;1x2,)+T(3;6),+6,-7,-8,T(8;4)+T(8;9),T(16;16),T(16,19),-16x2,- 19 |
| 8 | 44 | XY | -1,-1, +6,+6,+T(8;9),T(16;5), |
| 9 | 45 | XY | Der(1)?T(1;15;1),+3,+6,+T(8;9),T(16;16),+18 |
| 10 | 46 | XY | -1,Del(1),+Del(3),+3,+T(5;15),+6,+T(8;9),+T(3;14),T(15;5),-15 |

Del: Deletion
Der: Derivation/duplication
Dic: Dicentric
T: Translocation
Der-T: Derived band from translocation
Dic-T: Dicentric with translocation
+1: One more chromosome
-1: One less chromosome

**Supplementary Table 3: Most frequent human melanoma mutations found in the four models cell lines.**

| Model | Mutation | Allele fraction in cell line |
|---|---|---|
| **M1** | Pten (I32N) | 0.4 |
| | Trp53 (F131L, V140G, L142V) | 0.15 / 0.17 / 0.18 |
| | Erbb4 (S75F) | 0.47 |
| | Sf3b1 (E585D) | 0.14 |
| **M2** | Gnaq (Q209L) | 0.5 |
| **M3** | Gna11 (Q209L) | 0.4 |
| | Trp53 (R172H) | 1.0 |
| **M4** | Gnaq (Q209L) | 0.46 |
| | Gnaz (G230A) | 0.45 |
| | Kras (G12D) | 0.33 |
| | Sf3b1 (Q323H) | 0.41 |
| | Erbb4 (Y1250H) | 0.47 |

**Supplementary Table 4: Gate strategy for myeloid and lymphoid populations obtained by high parametric flow cytometry.**

| Immune Population | Markers |
|---|---|
| Myeloid cells: from Singlets Live $CD45^{+}$; Lin ($CD3^{-}$ $NK1.1^{-}$ $TCRb^{-}$ $TCR\gamma\delta^{-}$ $CD19^{-}$ $Ter119^{-}$) | |
| Macrophages | CD11b+ CD68hi CD64+ F480+ Ly6C-Ly6G-SiglecF- CD24- |
| Monocytes | CD11b+ CD68low CD64+ F480+ Ly6Chi Ly6G- SiglecF- CD24- |
| Neutrophils | CD11b+Ly6G+ Ly6Cint SiglecF- CD24+ |
| Eosinophils | CD11b+ Ly6G- Ly6Cint SiglecF+ F480+ CD64- CD24+ |
| Dendritic cells | Ly6G- SiglecF- CD11c+ MHC II+ CD64- CD24int/hi CD135+ |
| Lymphoid cells: all from Singlets Live CD45+ Lin (Ly6G- F480- CD19-) | |
| NK cells | CD3- NK1.1+ |
| T cells | CD3+ NK1.1- |
| CD4+ Tcells | CD3+ NK1.1- TCRb+ CD4+ |
| CD8+ Tcells | CD3+ NK1.1- TCRb+ CD8+ |
| gdTcells | CD3+ NK1.1- TCRgd+ TCRb- CD4- CD8- |
| Tregs | CD3+ NK1.1- TCRb+ CD4+ CD25+ Foxp3+ |
| DN T cells | CD3+ NK1.1- TCRb+ CD4- CD8- |

**Supplementary Table 7: Melanocytic Plasticity Signature (MPS) gene list.**

| Sign in the signature | Gene Symbol | Entrez Gene Name | Location |
|---|---|---|---|
| 1 | AKR1C3 | aldo-keto reductase family 1 member C3 | Cytoplasm |
| 1 | BMP1 | bone morphogenetic protein 1 | Extracellular Space |
| 1 | CRTAC1 | cartilage acidic protein 1 | Extracellular Space |
| 1 | ECEL1 | endothelin converting enzyme like 1 | Plasma Membrane |
| 1 | ERC2 | ELKS/RAB6-interacting/CAST family member 2 | Cytoplasm |
| 1 | FAM110C | family with sequence similarity 110 member C | Cytoplasm |
| 1 | FUT9 | fucosyltransferase 9 | Cytoplasm |
| 1 | GABRA2 | gamma-aminobutyric acid type A receptor alpha2 subunit | Plasma Membrane |
| 1 | GAP43 | growth associated protein 43 | Plasma Membrane |
| 1 | GREM1 | gremlin 1, DAN family BMP antagonist | Extracellular Space |
| 1 | HECW1 | HECT, C2 and WW domain containing E3 ubiquitin protein ligase 1 | Cytoplasm |
| 1 | KLHL1 | kelch like family member 1 | Cytoplasm |
| 1 | KRT12 | keratin 12 | Cytoplasm |
| 1 | LHFPL4 | LHFPL tetraspan subfamily member 4 | Other |
| 1 | NEFL | neurofilament light | Cytoplasm |
| 1 | NEFM | neurofilament medium | Plasma Membrane |
| 1 | NETO1 | neuropilin and tolloid like 1 | Extracellular Space |
| 1 | NKX2-2 | NK2 homeobox 2 | Nucleus |
| 1 | NSG2 | neuronal vesicle trafficking associated 2 | Cytoplasm |
| 1 | OCIAD2 | OCIA domain containing 2 | Cytoplasm |
| 1 | OTOP1 | otopetrin 1 | Plasma Membrane |
| 1 | PDE3B | phosphodiesterase 3B | Cytoplasm |
| 1 | PTPRN2 | protein tyrosine phosphatase, receptor type N2 | Plasma Membrane |
| 1 | PTPRT | protein tyrosine phosphatase, receptor type T | Plasma Membrane |
| 1 | SIGLEC15 | sialic acid binding Ig like lectin 15 | Plasma Membrane |
| 1 | SLC13A5 | solute carrier family 13 member 5 | Plasma Membrane |
| 1 | SLC9A2 | solute carrier family 9 member A2 | Plasma Membrane |
| 1 | SLITRK6 | SLIT and NTRK like family member 6 | Plasma Membrane |
| 1 | SNAP91 | synaptosome associated protein 91 | Plasma Membrane |
| 1 | STON2 | stonin 2 | Cytoplasm |
| 1 | TAC1 | tachykinin precursor 1 | Extracellular Space |
| 1 | VAT1L | vesicle amine transport 1 like | Other |
| 1 | WNT5A | Wnt family member 5A | Extracellular Space |
| -1 | ALX1 | ALX homeobox 1 | Nucleus |
| -1 | BRD7 | bromodomain containing 7 | Nucleus |
| -1 | DTD1 | D-tyrosyl-tRNA deacylase 1 | Cytoplasm |
| -1 | GRSF1 | G-rich RNA sequence binding factor 1 | Cytoplasm |
| -1 | HCN1 | hyperpolarization activated cyclic nucleotide gated potassium channel 1 | Plasma Membrane |
| -1 | LTA4H | leukotriene A4 hydrolase | Cytoplasm |
| -1 | OXCT1 | 3-oxoacid CoA-transferase 1 | Cytoplasm |
| -1 | PATJ | PATJ, crumbs cell polarity complex component | Plasma Membrane |
| -1 | PLXNC1 | plexin C1 | Plasma Membrane |
| -1 | SSBP4 | single stranded DNA binding protein 4 | Nucleus |
| -1 | TELO2 | telomere maintenance 2 | Cytoplasm |
| -1 | TMEM177 | transmembrane protein 177 | Cytoplasm |

##### Supplementary Table 8: Single variable Cox-PH model statistics for MPS and TIDE predictors

| Survival | Predictor | HR | CI95% | P-value |
|---|---|---|---|---|
| **Van Allen data set[4*]** | | | | |
| OS | MPS | 2.72 | 1.15-6.44 | 2.26E-02 |
| OS | TIDE | 5.04 | 2.19-11.62 | 1.45E-04 |
| PFS | MPS | 2.88 | 1.36-6.11 | 5.86E-03 |
| PFS | TIDE | 3.49 | 1.69-7.2 | 7.21E-04 |
| **Hugo-Riaz data set[8,19]** | | | | |
| OS | MPS | 2.17 | 1.02-4.6 | 4.38E-02 |
| OS | TIDE | 1.97 | 0.92-4.21 | 8.12E-02 |

*The predictors in this data set were adjusted by the age. HR: Hazard ratio

##### Supplementary Table 9: Multivariable Cox-PH model statistics for MPS and TIDE

| Survival | Variable | HR | CI95% | P-value |
|---|---|---|---|---|
| **Van Allen data set[4*]** | | | | |
| OS | MPS | 2.27 | 0.95-5.43 | 6.67E-02 |
| OS | TIDE | 4.7 | 1.99-11.12 | 4.24E-04 |
| PFS | MPS | 2.96 | 1.35-6.51 | 6.95E-03 |
| PFS | TIDE | 3.6 | 1.68-7.68 | 9.44E-04 |
| **Hugo-Riaz data set[8,19]** | | | | |
| OS | MPS | 2.25 | 1.06-4.8 | 3.49E-02 |
| OS | TIDE | 2.05 | 0.96-4.4 | 6.49E-02 |

*The predictors in this data set were adjusted by the age. HR: Hazard ratio

**Supplementary Tables provided upon request:**

**Supplementary Table 1. Nonsynonymous single nucleotide variants (SNVs) obtained from whole-exome sequencing of the GDAs and cell lines from the four models.**

**Supplementary Table 5. Gene expression in the untreated cell line-derived allografts from the four melanoma models.**

**Supplementary Table 6. Pairwise Gene Set Enrichment Analysis (GSEA) of the untreated cell line-derived allografts from the four models.**
